# DNA Wrapping by a Tetrameric Bacterial Histone

**DOI:** 10.1101/2025.05.08.652872

**Authors:** Yimin Hu, Samuel Schwab, Kaiyu Qiu, Yunsen Zhang, Kerstin Bär, Heidi Reichle, Aurora Panzera, Andrei N. Lupas, Marcus D. Hartmann, Remus T. Dame, Vikram Alva, Birte Hernandez Alvarez

## Abstract

Histones are conserved DNA-packaging proteins found across all domains of life. In eukaryotes, canonical histones form octamers that wrap ∼147 base pairs of DNA into nucleosomes — the fundamental units of chromatin. In archaea, histones form dimers that further multimerize into extended hypernucleosomes along DNA. Although bacteria were long thought to lack histones, recent studies have uncovered histone homologs in diverse bacterial lineages, many of which possess key DNA-binding features. We previously characterized HBb, a bacterial histone from *Bdellovibrio bacteriovorus*, which binds DNA as a dimer and induces bending. Here, we describe HLp from *Leptospira perolatii*, a representative of a distinct bacterial histone group. Crystallographic and biophysical analyses reveal that HLp forms stable tetramers. Like HBb, HLp binds DNA non-specifically; however, it adopts a different mode of interaction — wrapping ∼60 base pairs of DNA around its tetrameric core. This wrapping mode, supported by molecular dynamics simulations and DNA-binding assays, promotes DNA compaction and alters its topology. When expressed heterologously in *Escherichia coli*, HLp reorganizes nucleoid morphology, consistent with a role in chromatin organization. These findings expand the known repertoire of histone-DNA interaction in bacteria and underscore the structural and functional diversity of histone-based genome organization across the tree of life.

## Introduction

Histones are DNA-organizing proteins conserved across all domains of life. They share a characteristic histone fold — comprising three α-helices (α1, α2, α3) connected by two loops — that mediates histone-histone and histone-DNA interactions^1, 2^. In eukaryotes, histones assemble into nucleosomes, which serve as the basic units of chromatin. Nucleosomes consist of an octamer, composed of two copies each of histones H2A, H2B, H3, and H4, around which ∼147 base pairs (bp) of DNA wraps^1, 2,3^. These nucleosomes are arranged like beads on a string and are stabilized by linker histones (H1, H5), which bind DNA between adjacent nucleosomes^4, 5^. A hallmark of eukaryotic histones is their unstructured N-terminal tails, which undergo extensive post-translational modifications to regulate chromatin structure and gene expression^6,7^.

Histones were long believed to be unique to eukaryotes, although they play a central role in chromatin packaging and regulation. However, homologs have since been identified in archaea and, more recently, in bacteria — revealing that histone-based DNA organization is an ancient and widespread strategy for genome compaction. Despite this shared ancestry, key differences exist across domains. While eukaryotic histones are highly conserved and structurally uniform, forming nucleosomes with defined stoichiometry, prokaryotic histones exhibit far greater structural and functional diversity. Notably, N-terminal tails are generally absent in prokaryotes; however, histones from Asgard archaea — the closest known prokaryotic relatives of eukaryotes — do possess such tails^8, 9, 10, 11^. This suggests that chromatin regulation via tail modifications may have emerged prior to the evolution of the eukaryotic nucleus^9^. In both archaea and bacteria, genome organization has traditionally been attributed to nucleoid-associated proteins (NAPs), such as HU, H-NS, and IHF, which lack the histone fold but perform analogous roles in DNA compaction and transcriptional regulation^12, 13, 14, 15^. Based on sequence analysis and structural predictions, prokaryotic histones are broadly categorized into two major families: (i) nucleosomal histones, found exclusively in archaea, and (ii) α3 histones, present in both archaea and bacteria^9^. The α3 histones are defined by a shorter α2 helix and a truncated α3 helix, and exhibit extensive variation in their quaternary structures and domain organizations, leading to classification into multiple subfamilies^9^.

Archaeal histones are more abundant and better characterized than their bacterial counterparts, particularly the nucleosomal variants. A well-established example is HMfB from *Methanothermus fervidus*, which forms homodimers that tetramerize upon DNA binding, assembling into continuous helices where each dimer binds approximately 30 bp of DNA^16, 17^. Similarly, HTkA from *Thermococcus kodakarensis* has been shown to influence chromatin compaction and significantly modulate transcription initiation and elongation^18^. Beyond the nucleosomal type, non-nucleosomal archaeal histones exhibit even greater structural diversity. Members of the face-to-face (FtF) histone subfamily — part of the α3 histone group — form homotetramers that are predicted to wrap DNA^9^. This architecture is fundamentally distinct from the spiraling assembly seen in nucleosomal histones, as demonstrated by the crystal structure of HTkC from *T. kodakarensis*^9^. Another example, MJ1647 from *Methanocaldococcus jannaschii*, belongs to a histone subgroup found exclusively in *Methanococcales*. Unlike canonical histones, MJ1647 tetramerizes through its C-terminal helices rather than the histone fold and binds DNA in a bridging rather than wrapping mode^19^.

Bacterial histones are rare, present in fewer than 2% of sequenced bacterial genomes, in stark contrast to the NAP HU, which is found in over 90% of genomes^20^. Their limited distribution, relatively recent discovery, and frequent occurrence in uncultured or metagenomically characterized bacteria have left most bacterial histones uncharacterized at the functional level^21^. Among those identified, two main families dominate, α3 histones and DUF1931 pseudodimeric histones, both of which also occur in archaea^9^. The DUF1931 proteins, exemplified by structures from *Thermus thermophilus* (PDB: 1WWI) and *Aquifex aeolicus* (PDB: 1R4V), form pseudodimers but lack the conserved residues required for DNA binding and nucleosome-like assembly^9, 22, 23^. In contrast, α3 histones possess DNA-binding residues and are subdivided into five distinct subfamilies: (i) FtF histones, (ii) bacterial dimer histones, (iii) ZZ histones, (iv) phage histones, and (v) Rab GTPase histones^9^. The recent characterization of HBb from *Bdellovibrio bacteriovorus*, a member of the bacterial dimer subfamily, provided the first experimental evidence of a functional bacterial histone. HBb is essential for viability and binds genomic DNA non-specifically as a dimer^20, 24^. Although initially thought to bind without affecting DNA topology, structural and biophysical analyses later revealed that HBb induces local DNA bending, employing an interaction mode reminiscent of eukaryotic histones^24^.

To broaden our understanding of bacterial histone diversity, we investigated HLp from *Leptospira perolatii*, a member of the FtF subfamily of α3 histones. Crystallographic and biophysical analyses show that HLp forms stable tetramers. Structures of HLp in both free and DNA-bound forms reveal extensive protein-DNA contacts and a tetrameric architecture resembling that of archaeal HTkC^9^. Remarkably, HLp wraps ∼60 bp of DNA around its core, representing the first example of DNA wrapping by a bacterial histone. This mode of interaction, corroborated by molecular dynamics simulations and DNA-binding assays, drives DNA compaction and topological change, extending the known mechanisms of histone-DNA association in bacteria.

## Materials and methods

### Bioinformatic analyses

For cluster analysis, we used the histone sequences HMfB from *M. fervidus* (UniProtKB ID: P19267), HBb from *B. bacteriovorus* (Q6MRM1), the *A. aeolicus* pseudodimeric histone (O66665), the FtF histone from *L. interrogans* (Q8F3E8), and HTkC from *T. kodakarensis* (Q5JDW7) as queries in BLAST searches to retrieve archaeal and bacterial histone homologs. Searches were performed via the NCBI BLAST webserver with the ‘Max target sequences’ parameter set to 5000^25, 26^. Sequences shorter than 50 residues, longer than 200, or annotated as ‘partial’ or ‘fragment’ were excluded to eliminate truncated entries and multidomain proteins. The full-length hits were pooled and filtered using MMseqs2 to reduce redundancy at 80% pairwise sequence identity over 80% length coverage^27^. The resulting non-redundant dataset of 3341 sequences was clustered using CLANS based on all-against-all pairwise sequence similarities computed via BLAST^28, 29^. Clustering was performed to equilibrium using an E-value cut-off of 1e-6. Genome neighborhood analysis was carried out using the EFI Genome Neighborhood Tool, and structural predictions were generated using the AlphaFold2 (AF2)^30, 31^.

### Bacterial strains and cultivation

Cloning and plasmid amplification were conducted using *Escherichia coli* Top10. *E. coli* BL21(DE3) and *E. coli* Mutant56(DE3) were used as hosts for protein expression. Bacteria were grown in LB media supplemented with appropriate antibiotics.

### Cloning, plasmids, and synthetic DNA

The nucleotide sequences of the HLp-encoding gene CH373_10155 [GenBank: NPDY01000005.1, 141608 – 141799 (+)] from *L. perolatii strain FH1-B-C1* and the *hmfB* gene (GenBank: M34778.1) were codon-optimized and synthesized (BioCat GmbH) for protein expression in *E. coli* (Fig. S1A).

HLp was expressed fused to an N-terminal (histidine)6 tag using the vector pET- 30a(+), and HMfB was expressed using pET-28a(+). Ubiquitin from *Caldiarchaeum subterraneum* (*Cs*Ub), used as a control for light microscopy imaging, was expressed from pET-30b(+)^32^.

Synthetic oligonucleotides (Merck) used in this study are listed in Table S1. For DNA binding analyses, complementary single-stranded (ss) oligonucleotides were mixed, heated to 95 °C for 5 min, and slowly cooled to room temperature to generate double-stranded (ds) DNA fragments. The 600-bp-GC40 and 240-bp-GC40 DNA fragments used in DNA binding assays were amplified by PCR using the vector pETHis1a as the template^33^.

### Protein expression and purification

HMfB and HLp were expressed in *E. coli* Mutant56(DE3) and *E. coli* BL21(DE3), respectively. Bacterial cultures were maintained in LB broth supplemented with kanamycin at 37 °C. At an optical density of OD600 = 0.6, isopropyl-β-d-thiogalactoside (IPTG) was added at a final concentration of 1 mM to induce protein expression. Cells were agitated at 37 °C for an additional 4 h. Cells were pelleted and resuspended in lysis buffer containing 20 mM Tris, pH 8.0, 300 mM NaCl, and 10 mM imidazole, supplemented with protease inhibitor mix (cOmplete™, EDTA-free Protease Inhibitor Cocktail, Roche), 0.1 mM PMSF, 3 mM MgCl2 and DNase I (AppliChem GmbH). Cells were lysed using a French press, and cell debris and insoluble material were pelleted by centrifugation at 95,000 x g for 45 min. The supernatant was filtered through a 0.45 μm filter and applied to a 5 mL HisTrap column (Cytiva). Bound proteins were eluted using a linear imidazole gradient, ranging from 10-500 mM, in the aforementioned buffer. For HLp, the (histidine)6 tag was cleaved with *Tobacco Etch Virus* (TEV) protease. A second purification step using a HisTrap column was performed using the aforementioned buffers containing 150 mM NaCl to separate the protein from the cleaved tag. Purified proteins were dialyzed against 20 mM Tris buffer at the indicated pH with 150 mM NaCl and concentrated using an Amicon Ultra Centrifugal Filter Device (3 kDa MWCO, Millipore). Protein purity was confirmed by SDS-PAGE (mPAGE 4-12% Bis-Tris Precast Gel, Millipore), and protein concentration was determined spectrophotometrically or using a BCA protein assay (ThermoFisher Scientific). For the tethered particle motion (TPM) assay, purified HLp was dialyzed against 50 mM Tris, pH 7.0, 75 mM KCl, and 10% glycerol.

### Circular dichroism (CD) spectroscopy

HLp was dialyzed against 10 mM phosphate buffer, pH 8.0, 75 mM KF, and diluted to a final concentration of 15 μM. CD spectra were recorded on a Jasco J-810 spectrometer (JASCO) in the wavelength range of 190-250 nm using a cuvette with a 1 mm path length and a 100 nm/min reading speed. A total of ten single spectra were recorded and averaged. To analyze the thermal stability of HLp, the ellipticity (θ) was measured at 222 nm over a temperature gradient of 10-100 °C, using a ramp of 1 °C/min, a data pitch of 0.5, and a response time of 1 s. Data analysis, blank subtraction, and curve smoothing were performed using the Spectra Manager software suite (JASCO). CD spectra and thermal melting curves were plotted using GraphPad Prism 9 (GraphPad Software, Inc.).

### Size exclusion chromatography coupled with multi-angle light scattering (SEC-MALS)

The HLp-DNA complex was analyzed by SEC-MALS following incubation of HLp with the 30-bp-GC40 DNA fragment at a protein-to-DNA molar ratio of 2:1. All samples were incubated at 37 °C for 10 min, and aggregates were removed by centrifugation. Samples were applied onto a Superdex 75 Increase 10/300 GL column (Cytiva), pre-equilibrated with SEC buffer (25 mM Tris, 50 mM NaCl, 50 mM KCl, pH 7.5). The run was performed at a 0.5 mL/min flow rate on a 1260 Infinity II HPLC system (Agilent) coupled to a miniDAWN TREOS and Optilab T-rEX refractive index detector (Wyatt Technology). HLp and the 30 bp-GC40 DNA fragment were detected at 215 nm and 260 nm, respectively. Measurements were performed in triplicate, and molecular mass distributions were calculated using the ASTRA v.7.3.0.18 software suite (Wyatt Technology).

### Crystallization, data collection, and structure determination

Crystallization trials were set up by mixing 300 nL of protein with 300 nL of reservoir solution in 96-well sitting-drop vapor-diffusion plates using commercially available screens with reservoir volumes of 100 μL. Without DNA, HLp was prepared at 11 mg/mL in 20 mM Tris, pH 8.0, and 150 mM NaCl. The best diffracting crystals were obtained with a reservoir solution containing 0.1 M citric acid, pH 5.0, and 3.15 M ammonium sulfate. For co-crystallization of HLp and DNA, 500 µM HLp was mixed with 500 µM 30-bp-GC40 dsDNA (Table S1) and incubated at 37 °C for 10 min, after which aggregates were removed by centrifugation. The conditions under which the crystals used for structure determination grew were 0.1 M HEPES, pH 7.5 and 25% PEG 6000 for HLp-DNA_1, and the Morpheus condition G12 [0.1 M carboxylic acids, 0.1 M Morpheus Buffer System 3, pH 8.5 and 50% (V/V) Morpheus Precipitant Mix 4] for HLp-DNA_2. For cryo-protection, the crystals were transferred to droplets of their reservoir solution spiked with 30% glycerol (free HLp) or 15% PEG 400 (HLp-DNA_1), loop-mounted, and flash frozen in liquid nitrogen. Data were collected at beamline X10SA of the Swiss Light Source (Villigen, Switzerland) at 100 K, using an EIGER X 16M hybrid pixel detector (Dectris, Ltd.). Data were reduced, processed, and scaled using XDS^34^. Due to pronounced anisotropy, diffraction data for HLp-DNA_2 were submitted to the STARANISO server for ellipsoidal truncation and anisotropic scaling, following the unmerged data protocol^35^.

The structure of free HLp was solved by molecular replacement (MR) using MOLREP and an AlphaFold2 prediction as a search model, locating one HLp dimer in the asymmetric unit (ASU)^31, 36, 37^. The DNA-bound structures were solved by MR using MOLREP and the refined free HLp coordinates as a search model, locating an HLp dimer and 16 bp of dsDNA in the ASU for HLp-DNA_1, and a monomer and 15 nucleotides of ssDNA in the ASU for HLp-DNA_2. The three structures were modeled, refined, and finalized in cycles of manual modeling in Coot and refinement with REFMAC5^38, 39^. Data processing and refinement statistics are given in Table S3. The coordinates and structure factors have been deposited in the PDB under the accession numbers 9QT0 (free HLp), 9QT1 (HLp-DNA_1), and 9QT2 (HLp-DNA_2). All structures were visualized using PyMOL (The PyMOL Molecular Graphics System, Version 3.0 Schrödinger, LLC.).

### Molecular dynamics (MD) simulation

We performed all-atom molecular dynamics (MD) simulations in GROMACS (version 2023.2) to study two possible topologies of the HLp-DNA complexes formed by bridging or wrapping^40^. The DNAs in the HLp-DNA_1 and HLp-DNA_2 structures were extended through crystal symmetry. The obtained structures were superimposed based on the central HLp, the excess DNA fragments were trimmed up to the overlap point, and then the remaining DNA fragments were connected to obtain the two initial structures of HLp-DNA complexes (Fig. 5 and Fig. S6). Two initial models were constructed for each binding mode (wrapping and bridging) based on the HLp-DNA_1 and HLp-DNA_2 structures. One representative model for each binding mode was pursued further. The constructed models were then preprocessed using PDBFixer (version 1.10, https://github.com/openmm/pdbfixer) to add missing hydrogen atoms and adjust chain configurations as necessary. The CHARMM all-atom force field (CHARMM36m) was applied to parameterize and describe both intra- and intermolecular interactions between the protein and DNA^41^. Ion parameters and the TIP3P water model were selected following the CHARMM36m recommendations^42^.

Each system was solvated in a sufficiently large cubic water box to ensure a minimum water layer of 1.5 nm surrounding the protein-DNA complex in all directions. Sodium and chloride ions were added to neutralize the system. Periodic boundary conditions (PBC) were imposed to eliminate edge effects, enabling the simulation to be conducted under an infinite solvent environment.

Each system underwent a multistage process of energy minimization and equilibration before production simulation. First, a harmonic positional restraint of 1000 kJ/(mol·nm²) was applied to the backbones of both proteins and DNAs to relax the distributions of water molecules and ions. The system was then subjected to 5000 steps of conjugate gradient and steepest descent minimization until the maximum force on any atom dropped below a predefined threshold of 1000 kJ/(mol·nm²). Following energy minimization, each complex system was equilibrated in multiple stages using the canonical (NVT) and isothermal-isobaric (NPT) ensembles. During the NVT phase, the system was gradually heated to 303.15 K with a V-rescale thermostat^43^. In the NPT phase, density equilibration was achieved at a constant pressure of 1 atm using a Parrinello-Rahman barostat^44^. Once equilibration was complete, a 1 μs production simulation was conducted on the processed protein-DNA complex, with the integration carried out using the leap-frog algorithm at a timestep of 2 fs. Meanwhile, a 1.2 nm cutoff was set for van der Waals (VDW) and electrostatic potentials to calculate the short-range interaction, and the Particle Mesh Ewald (PME) method was employed to compute the long-range interaction^45^. To enhance computational efficiency, the LINCS algorithm was employed to constrain hydrogen bond lengths in water molecules and the backbones of both proteins and DNAs^46^. Two independent 1 μs simulations were performed for each system to ensure the robustness of the results (SI movie 1 and 2). Both simulations exhibited highly similar behaviors in either the wrapping or the bridging model; therefore, the analysis was conducted on one representative trajectory for each system. To assess global structural stability, we calculated the root mean square deviation (RMSD) of the DNA backbone relative to the initial structure. Binding fluctuations were evaluated by determining the difference between the contact number of each frame and the average contact number over all captured frames. Protein-DNA contacts were defined by the presence of any protein-heavy atom within 0.4 nm of a DNA-heavy atom. Hydrogen bonds were identified with a distance cutoff of 0.3 nm for the donor-to-acceptor (D-A) distance and an angular cutoff of 150° for the donor-hydrogen-acceptor (D-H-A) angle. To further study the binding dynamics of the local interaction sites, we calculated the binding contact number for each residue throughout the simulation and manually selected residues to define four binding sites (Fig. 5C and Table S4).

To further characterize and understand the binding dynamics of the proposed wrapping model, an unwrapping simulation was performed in the framework of a two-step steered molecular dynamics (SMD)^47^, with the constructed wrapping model (Fig. S6) as the starting structure. The first SMD simulation targeted the dissociation of the B-site 2, followed by a second SMD simulation that used the last snapshot from the B-site dissociation as the starting structure to investigate the unbinding of the A-site. In the SMD setup, a reaction coordinate was defined by connecting the center of mass of the relevant DNA regions (residues 1 to 12) to that of the protein, ensuring that the applied force was directed along an axis expected to promote the unwrapping process. A pulling velocity of 0.002 nm/ps and a harmonic spring constant of 500 kJ/(mol·nm²) were selected to balance the computational feasibility with the need to capture quasi-equilibrium behavior. During the production simulation of SMD, force and extension data were recorded at regular intervals to allow integration of the force over distance and estimation of the work required for unwrapping. Simultaneously, the temporal evolution of protein-DNA contacts was monitored using a 0.4 nm cutoff to pinpoint specific events where the DNA disengaged from the binding interface. Once both B-site 2 and A-site 2 were dissociated, we integrated the force-distance curves obtained from the SMD simulation to generate dissociation energy profiles. All simulations were analyzed using VMD (version 1.9.4) and the MDAnalysis package (version 2.8.0), with Matplotlib and Seaborn employed for visualization^48, 49^.

### DNA binding assays

All DNA binding assays were performed as described previously, with minor modifications^24^.

#### Electrophoretic mobility shift assay (EMSA)

The 80-bp dsDNA fragment, 30-bp dsDNA fragments of various GC content (30%, 40%, 50%, and 60%), and the GeneRuler 1 kb Ladder (ThermoFisher Scientific) were used for *in vitro* DNA binding tests. For comparative DNA binding analysis, the proteins were mixed with the annealed dsDNA fragments at the indicated molar ratios in a binding buffer containing 25 mM Tris, pH 8.0, 50 mM NaCl, and 50 mM KCl. Following incubation at 37 °C for 10 min, the samples were separated on a 6% DNA retardation gel (ThermoFisher Scientific). To analyze the binding of HLp and HMfB to the GeneRuler 1 kb Ladder, the proteins and the DNA were mixed at the indicated mass ratios in the binding buffer and incubated at 37 °C for 10 min. Samples were separated on a 1% agarose gel, and the DNA was visualized using SYBR Gold nucleic acid gel stain (ThermoFisher Scientific) and imaged using the Fusion SL imaging system (Vilber).

#### Micrococcal nuclease (MNase) digestion assay

The 600-bp-GC40 DNA fragment was amplified using primers pET-600-bp-GC40-F and pET-600-bp-GC40-R with pETHis1a as a template, followed by purification using the QIAquick PCR Purification Kit (QIAGEN). For HLp-DNA complex formation, 720 ng of HLp was mixed with 900 ng of 600-bp-GC40 DNA in MNase buffer (10 mM Tris, pH 8.0, 75 mM NaCl) and incubated at 37 °C for 10 min. Indicated amounts of MNase (New England Biolabs) were added, and digestion was performed in a reaction volume of 80 μL at 37 °C for 10 min. As a positive control, HMfB (540 ng) was preincubated with 600-bp-GC40 DNA (900 ng) and treated with the indicated amounts of MNase at 37 °C for 15 min. EDTA and SDS were added to final concentrations of 95 mM and 0.5%, respectively, to stop the reaction. DNA fragments protected by bound proteins were purified by phenol/chloroform extraction followed by ethanol precipitation. The purified DNA fraction was dissolved in 5 μL of water and separated on a 10% Novex TBE gel (ThermoFisher Scientific). The gel was stained with SYBR Gold nucleic acid gel stain (ThermoFisher Scientific) and imaged using the Gel Doc XR+ imaging system (Bio-Rad Laboratories). All experiments were performed at least in triplicate, and the length of the selected DNA bands was calculated using Image Lab 6.1 (Bio-Rad Laboratories). For better visualization, the image background was subtracted using the rolling-ball algorithm with a radius of 50 pixels in FIJI (version 2.3.0, https://fiji.sc/)^50^.

#### Tethered particle motion (TPM) assay

TPM experiments were performed as previously described in 50 mM Tris, pH 7.0, and 75 mM KCl ^51^. A standard deviation cutoff of 8% and an anisotropic ratio cutoff of 1.3 were used to select single-tethered beads^51^. Measurements at each HLp concentration were done in triplicate. Means and standard deviations of the individual measurement series for each HLp concentration were calculated by maximum likelihood estimation, assuming a normal distribution. Outliers with a robust Z-score >3 or <-3 were not considered for fitting. The “line to guide the eye” was generated by fitting the means to a logistic function. A custom Python script was used to fit and plot the TPM data. For plotting, the means of the three individual measurements were averaged for each measured concentration, and the standard deviations were error-propagated 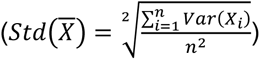.

#### DNA topology assay

Plasmid pUC19 was purified from *E. coli* Top10 using NucleoSpin Plasmid QuickPure Kit (Macherey-Nagel). For topological relaxation, the plasmid pUC19 was treated with the nicking endonuclease Nb.BsrDI (New England BioLabs), followed by ligation with T4 DNA ligase (ThermoFisher Scientific). Plasmid relaxation was verified by agarose gel electrophoresis. 200 ng of relaxed pUC19 was mixed with HLp or HMfB at indicated protein-to-DNA mass ratios in assay buffer containing 10 mM Tris, pH 7.0, and 75 mM NaCl) and incubated at room temperature for 30 min. Then, 1 U of Topoisomerase I (ThermoFisher Scientific) was added, and the reaction mixture was incubated at 37 °C for 30 min in a final volume of 50 μL. DNA was purified by phenol/chloroform extraction followed by ethanol precipitation. Purified DNA samples were separated on a 0.8% TAE agarose gel. Plasmids were visualized by staining with SYBR Gold nucleic acid gel stain (ThermoFisher Scientific) and imaged using the Gel Doc XR+ imaging system (Bio-Rad Laboratories).

#### Ligase-mediated circularization assay

The 240-bp-GC40 DNA fragment was amplified from plasmid pETHis1a using primers pET-240-bp-GC40-Fp and pET-240-bp-GC40-Rp (Table S1) and purified using QIAquick PCR Purification Kit (QIAGEN)^33^. 400 ng of DNA was incubated with HMfB and HLp at the indicated protein-to-DNA mass ratios in LMC buffer (10 mM Tris, pH 7.0, 75 mM KCl, and 5% glycerol) in a total reaction volume of 20 μL at room temperature for 30 min. T4 DNA ligase buffer and T4 ligase at a final concentration of 1 U/μL were added in a final reaction volume of 100 μL. A sample not treated with T4 DNA ligase was used as a control. The sample was incubated for 24 h at room temperature, and the DNA was purified by phenol/chloroform extraction and ethanol precipitation. Half of each sample was treated with 1 U of T5 exonuclease (New England BioLabs) in the appropriate buffer at 37 °C for 1 h. DNA samples were separated on a 2% TAE agarose gel, stained with SYBR Gold nucleic acid gel stain (ThermoFisher Scientific), and imaged using the Gel Doc XR+ imaging system (Bio-Rad Laboratories).

#### Microscale thermophoresis (MST)

To determine the DNA binding affinity of HLp to the Cy5 labeled 80-bp and 80-bp- GC40 dsDNAs, a dilution series of HLp was prepared in 20 mM Tris, pH 8.0, 150 mM NaCl, and titrated against the dsDNA fragments at a concentration of 10 nM. The HLp-DNA mixtures were incubated at 37 °C for 10 min and loaded into Monolith NT premium capillaries (MO-K025, NanoTemper Technologies) after centrifugation to remove precipitates. HMfB was used as a positive control and titrated against both DNA fragments at a concentration of 20 nM. HMfB-DNA mixtures were incubated at room temperature for 5 min, centrifuged, and loaded into Monolith NT capillaries (MO-K022, NanoTemper Technologies). All measurements were performed in triplicate at 25 °C using the Monolith NT 115 instrument with a Nano RED detector and MST power set to medium. MST data were analyzed by fitting them to a Kd model using MO Control (NanoTemper Technologies).

### Light microscopy imaging and data processing

Vectors pET-30a(+) and pET-30a(+) encoding *C. subterraneum* ubiquitin (*Cs*Ub) were transformed in *E. coli* BL21(DE3)^32^. The transformed strains, as well as non-transformed *E. coli* BL21(DE3) and *E. coli* Mutant56(DE3), served as negative controls.

#### Sample preparation

Overnight cultures of each strain were diluted 30-fold (V/V) in LB medium supplemented with the appropriate antibiotic and incubated at 37 °C, 170 rpm. Except for the negative controls, protein expression was induced with IPTG at a final concentration of 1 mM at an OD600 = 0.4-0.6. Following cultivation for another 4 h, 500 µL bacterial culture was pelleted. The cells were washed with PBS and fixed using 4% (V/V) formaldehyde. Fixation was stopped with glycine at a final concentration of 150 mM. Following two wash steps with PBS, the cells were incubated with 1 μg/mL DAPI solution (Invitrogen), washed, and resuspended in PBS. The treated cells were dropped onto a 2% agarose pad, which was placed upside down in a 35 mm imaging dish (High Glass Bottom, Ibidi) for imaging. Triplicates of each sample were analyzed.

#### Imaging and image processing

Light microscopy imaging was conducted using a Zeiss LSM 780 inverted confocal microscope (Carl Zeiss AG, Oberkochen, Germany) equipped with a 63 x oil/1.4NA oil-immersion objective. The 405 nm diode laser line was used for DAPI excitation and brightfield imaging. Airyscan datasets were processed with Airyscan software to generate 32-bit images using pixel reassignment and Wiener filter-based deconvolution^52^. Brightfield images were used to segment individual bacterial cells via a custom-developed Macro in FIJI, which included background subtraction and auto-thresholding^50^. Segmentation results were manually curated to eliminate artefacts. For fluorescence analysis, a maximum intensity projection was first applied to the Airyscan z-stacks. The segmented bacterial outlines were overlaid onto the fluorescence channel, followed by a second segmentation step on the fluorescence channel to identify nucleoid regions using auto-thresholding.

## Results

### Bioinformatic analysis

To explore the diversity and evolutionary relationships of bacterial FtF histones and identify candidates for experimental characterization, we performed a comparative sequence analysis based on a curated dataset comprising α3 histones from the FtF and bacterial dimer subfamilies, archaeal nucleosomal histones, and pseudodimeric DUF1931-family histones. Only proteins consisting solely of the histone fold, without additional domains, were included. All-against-all pairwise sequence similarities were analyzed using CLANS, enabling visualization of the sequence space occupied by FtF histones and assessment of their relationships to other prokaryotic histone families. The resulting cluster map reveals a clear separation among major prokaryotic histone families (Fig. 1A). DUF1931 pseudodimeric histones form a distinct, compact cluster, well separated from canonical histones. Archaeal nucleosomal histones group into a large, coherent cluster, with a small subgroup of closely related sequences extending from its core, including the *Haloferax volcanii* pseudodimer HstA (HVO_0520, UniProt D4GS56). Archaeal FtF histones, bacterial FtF histones, and bacterial dimer histones form separate, well-defined clusters. The adjacency of the bacterial FtF cluster to the archaeal FtF cluster suggests potential shared structural features, such as tetramerization.

**Fig. 1.**
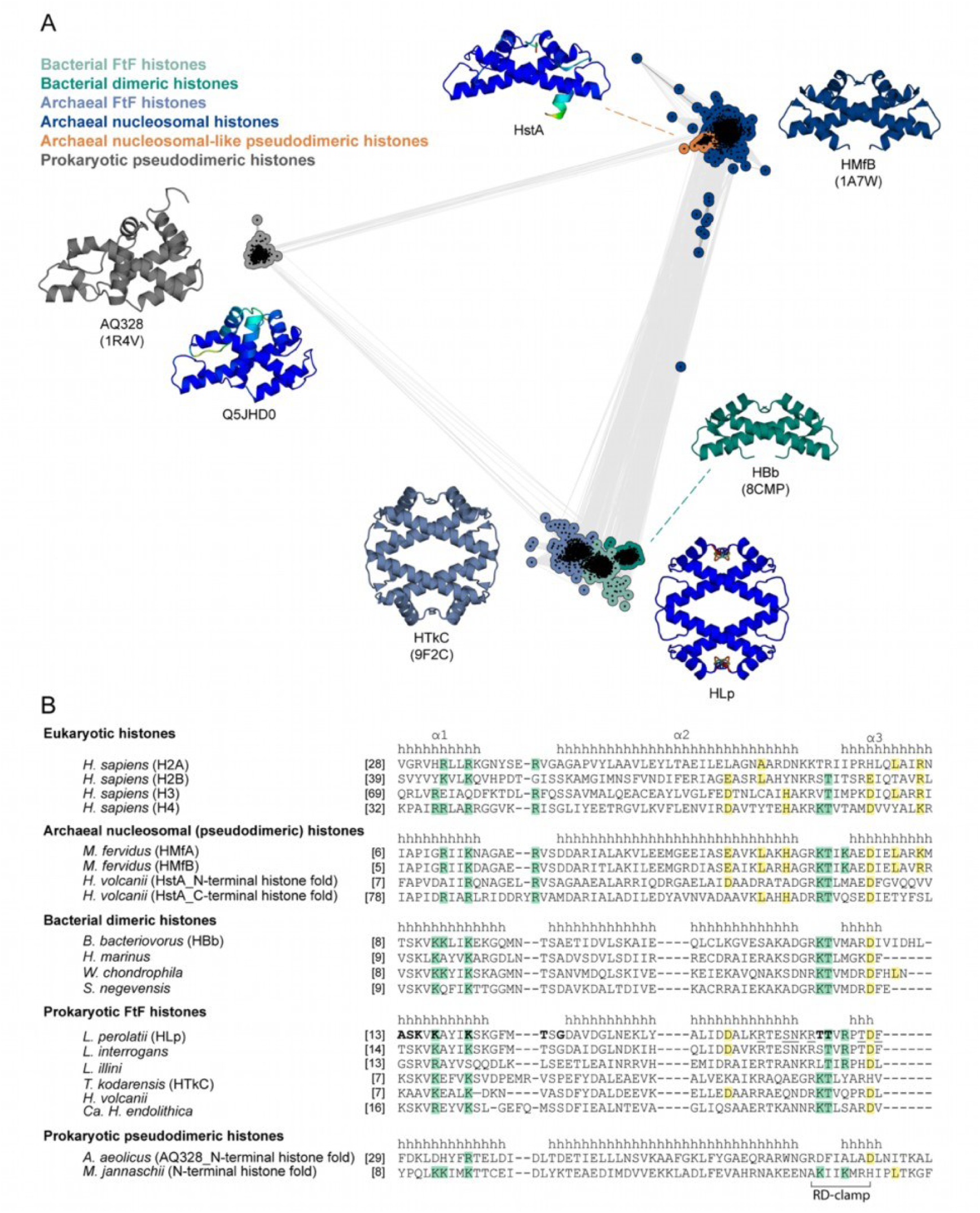
Comparative analysis of prokaryotic histones composed exclusively of histone fold domains. **A.** Cluster map of ∼3,300 prokaryotic histones containing only the histone fold domain. Each dot represents a single protein sequence, edges reflect pairwise sequence similarity. Structures of representative proteins — either crystal structures (PDB IDs in parentheses) or AlphaFold models — are shown in cartoon representation. AlphaFold models are colored by pLDDT confidence scores. Representative proteins include: HstA from *Haloferax volcanii* (D4GS56), an archaeal nucleosomal-like pseudodimeric histone; HMfB from *Methanothermus fervidus* (P19267), an archaeal nucleosomal histone; HBb from *Bdellovibrio bacteriovorus* (Q6MRM1), a bacterial dimeric histone; HLp from *Leptospira perolatii* (A0A2M9ZN55), a bacterial FtF histone; HTkC from *Thermococcus kodakarensis* (Q5JDW7), an archaeal FtF histone; and Q5JHD0 from *T. kodakarensis* (Q5JHD0) and AQ328 from *Aquifex aeolicus* (O66665), both prokaryotic pseudodimeric histones. **B.** Multiple sequence alignment of representative histones from both eukaryotes and prokaryotes, including *Homo sapiens* (H2A: P04908; H2B: P62807; H3: P68431; H4: P62805), *M. fervidus* (HMfA: P48781; HMfB: P19267), *H. volcanii* (HstA: D4GS56), *B.bacteriovorus* (HBb: Q6MRM1), *Hymenobacter marinus* (NCBI accession No.: ATH09486), *Waddlia chondrophila* (D6YWW1), *Simkania negevensis* (F8L7X8), *L. perolatii* (HLp: A0A2M9ZN55), *L. interrogans* (Q8F3E8), *Leptonema illini* (H2CFS2), *T. kodakarensis* (HTkC: Q5JDW7), *H. volcanii* (D4GZE0), Candidatus *Heimdallarchaeum endolithica* (A0A9Y1FPJ9), *A. aeolicus* (O66665), and *Methanocaldococcus jannaschii* (A0A832T4V6). α-helices are annotated with “h” based on crystal structures of H2A, HMfA, HBb, HLp, and AQ328. Conserved residues associated with DNA binding and oligomerization are highlighted in green and yellow, respectively. HLp residues contributing to tetramerization in the HLp crystal structure (PDB: 9QT0) are underlined, while those interacting with the DNA backbone in HLp-DNA complexes (PDB: 9QT1 and 9QT2) are shown in bold. Unless otherwise stated, accession numbers in parentheses refer to UniProtKB.

While FtF histones are widespread across archaeal phyla, in bacteria they are predominantly found in specific lineages, including Spirochaetota, Planctomycetota, Bdellovibrionota, and Myxococcota. The majority of these bacterial representatives are derived from uncultured organisms or metagenomic assemblies, limiting opportunities for functional characterization. Among the few exceptions, we identified two species with complete genomes from culturable organisms within the bacterial FtF cluster: *L. interrogans*, a pathogenic species and the primary causative agent of leptospirosis, and *L. perolatii*, a species occasionally associated with disease in humans and animals. The FtF histone from *L. interrogans* is known to be highly abundant and essential for viability, underscoring the functional relevance of this histone group^20^. Notably, neither *L. interrogans* nor *L. perolatii* encodes classical bacterial NAPs such as HU or Dps, suggesting that FtF histones may serve as the principal DNA organizers in these lineages. Consistent with this, we found only one histone homolog in each of the two *Leptospira* species. Interestingly, in many *Leptospira* species, the *hlp* gene is located adjacent to genes encoding the chromosome segregation protein SMC (WP_100713470.1), methionine aminopeptidase (MAP; WP_100713471.1), and an uncharacterized DUF350 domain-containing transmembrane protein (WP_100713472.1). This conserved genomic neighborhood hints at potential functional links between histone-mediated DNA organization and core aspects of cellular physiology. We focused our efforts on the histone homolog HLp from *L. perolatii*, thereby enabling the characterization of a previously unexplored bacterial FtF histone and advancing our understanding of histone-based chromatin organization in Leptospira, beyond the limited insights available from *L. interrogans*^53^. Moreover, AF structure predictions indicated that HLp forms a homotetramer, resembling that of the archaeal FtF histone HTkC from *T. kodakarensis*, further supporting its relevance as a representative of this bacterial subgroup^9^.

### Structure of HLp from *L. perolatii*

Based on the above bioinformatic analysis, HLp from *L. perolatii* was selected as a representative of the bacterial FtF histone group for structural and functional characterization. In sequence databases, two variants of the HLp protein are documented: a shorter form comprising 63 amino acid residues and a longer form with 77 residues, differing in their N-terminal regions (Fig. S1A). To determine the correct variant, we examined the genomic context of the *hlp* gene, particularly analyzing the position of the Shine-Dalgarno (SD) sequence — a ribosomal binding site crucial for initiating bacterial translation^54^. Our analysis revealed that the SD sequence aligns appropriately with the start codon of the shorter, 63-residue variant, suggesting that this form is the authentic translation product (Fig. S1A). HLp was recombinantly overexpressed in *E. coli* and purified to homogeneity, as confirmed by SDS-PAGE, which showed a single band at ∼10 kDa, consistent with its theoretical molecular weight of 7.1 kDa (Fig. S2A). SEC-MALS revealed a predominant species of 26 ± 0.3 kDa, indicating that HLp forms tetramers in solution (Fig. S2B). CD spectroscopy showed that HLp is a predominantly α-helical protein and unfolds upon heating, with a melting temperature of 57.1°C (Fig. S2C and D).

HLp crystallized under multiple conditions within a few days. The best data set was processed to a resolution of 1.30 Å, and the structure was solved by molecular replacement using an AF model as the search template (Fig. 2)^31, 36^. The entire chain, except for the first seven residues, was well resolved in the electron density map. The asymmetric unit contained an HLp dimer, which assembles into a homotetramer by crystallographic symmetry. This tetramer closely matches the AF model, with an RMSD value of 0.907 Å (Fig. S3A).

**Fig. 2.**
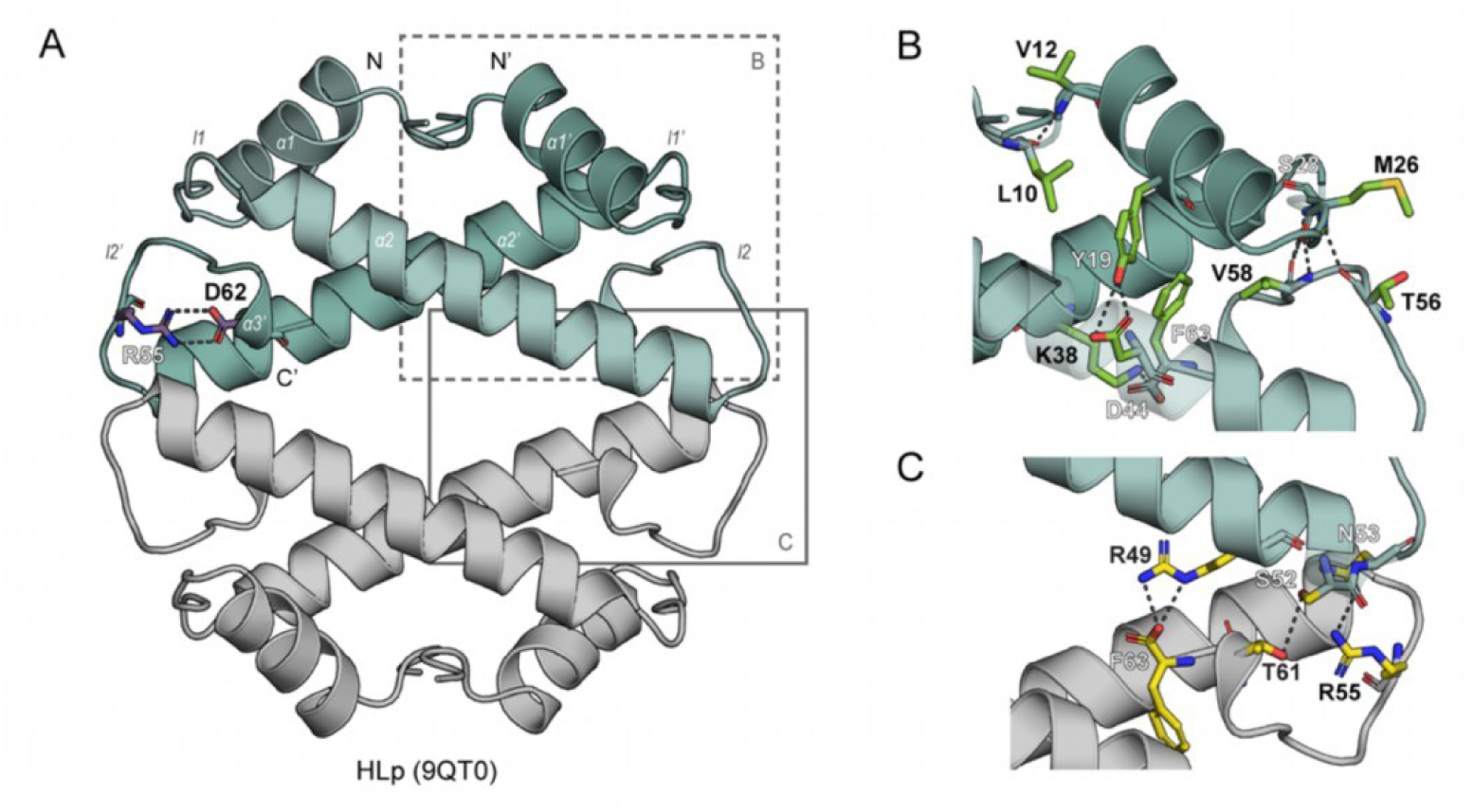
Crystal structure of the HLp tetramer. **A.** Crystal structure of the HLp tetramer (PDB: 9QT0) shown in cartoon representation. Residues R55 and D62, which form the RD clamp, are shown as sticks. Salt bridges are indicated as dashed lines. **B.** Close-up view of the HLp dimerization interface. Residues involved in dimerization are shown as green sticks, with hydrogen bonds depicted as dashed lines. **C.** Close-up view of the HLp tetramerization interface. Tetramerization-associated residues are shown as yellow sticks. Salt bridges and hydrogen bonds are indicated by dashed lines. In all panels, the contents of the asymmetric unit are shown in color, with selected symmetry mates in gray.

The HLp monomer exhibits the characteristic histone fold, comprising three α- helices (α1, α2, and α3) connected by two loops (l1 and l2) (Fig. 2A). The central α2 helix is one turn shorter than in the archaeal histone HMfB (Fig. S4), while the C-terminal α3 helix is truncated and forms a single helical turn composed of the final four residues — similar to what is observed in the bacterial dimeric histone HBb from *B. bacteriovorus*^24^. Within the dimer, the monomers are arranged in a head-to-tail orientation, with the interface formed primarily by the antiparallel crossing of the α2 helices at an angle of ∼40°. It is stabilized by hydrophobic interactions involving residues V16 and I20 (α1 helix), a leucine-rich stretch in the α2 helix (L35, L39, L42, and L47), and F63 (α3 helix), along with polar contacts between loop regions. Notably, M26 and S28 in l1 of one monomer interact with T56 and V58 in l2 of the opposing monomer (Fig. 2B).

Tetramerization is mediated by contacts between the C-terminal end of α2, loop l2, and α3 of opposing dimers. This includes a branched hydrogen-bonding network formed between monomers from opposing dimers: specifically, between R49, S52, and N53 of one, and F63, T61, and R55 of the other monomer, respectively (Fig. 2C). Notably, R49, N53, and R55 are conserved across predicted tetrameric bacterial FtF histones, suggesting a common mechanism of tetramer assembly (Fig. 1B). This mode of oligomerization is fundamentally distinct from that of archaeal nucleosomal histones such as HMfB, which assemble into extended spirals (Fig. S4B)^16^.

HLp also harbors a conserved RD-clamp — an intramolecular salt bridge between R55 and D62 — that stabilizes loop l2, a feature shared with both HMfB and HBb (Fig. 1B, Fig. 2A, and Fig. S4)^24, 55^. Compared to canonical histones, HLp shows partial conservation of residues implicated in DNA binding (Fig. 1B). Nevertheless, electrostatic surface potential analysis of the HLp tetramer reveals a continuous band of positive charge encircling the structure, consistent with a DNA-wrapping mode of interaction. (Fig. S3B)^56^.

### HLp binds non-specifically to DNA *in vitro*

We first assessed the DNA-binding ability of HLp using EMSAs. The 80-bp DNA fragment — previously shown to bind HMfB with high affinity — exhibited reduced mobility in polyacrylamide gels upon incubation with HLp (Fig. 3A), indicating the formation of HLp-DNA complexes^16, 57^. Complex formation was concentration-dependent, and the appearance of smeared bands suggested that the HLp-DNA complexes were less stable or more heterogeneous than those formed with HMfB.

**Fig. 3.**
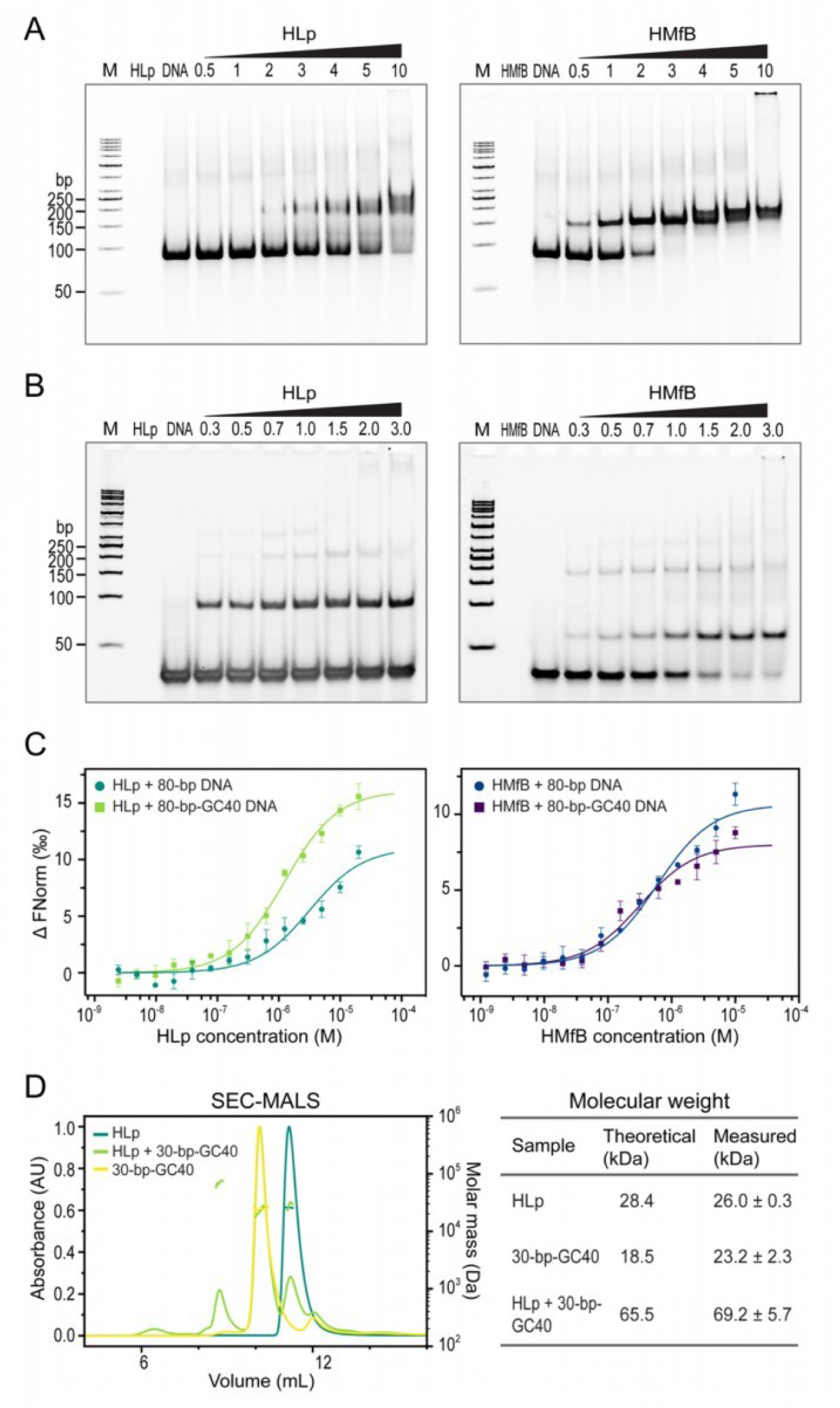
**HLp binds DNA *in vitro*.** EMSAs showing the binding of HLp and HMfB (control) to the 80-bp DNA fragment (**A**) and the 30-bp-GC40 DNA fragment (**B**). Increasing protein concentrations (lanes 4- 10), indicated as molar protein-to-DNA ratios, were incubated with the corresponding DNA and analyzed on a 6% polyacrylamide gel. **C.** MST measurements comparing the binding of HLp to the 80-bp and the 80-bp-GC40 DNA fragments, alongside HMfB. **D**. Profiles of SEC-MALS runs of HLp, the 30-bp-GC40 DNA, and a mixture of both; the table summarizes both theoretical and experimentally determined molecular weights.

To further investigate sequence preferences, we repeated the EMSA with shorter 30-bp DNA fragments varying in GC content. These assays yielded sharper band patterns, and HLp showed the highest affinity for DNA with 40% GC content (Fig. 3B and Fig. S5). Binding was confirmed by microscale thermophoresis (MST), which demonstrated that HLp interacts with DNA non-specifically, with the strongest binding to the DNA of 40% GC content and a dissociation constant in the low micromolar range (Fig. 3C and Table S2).

We next used size-exclusion chromatography coupled with SEC-MALS to determine the stoichiometry of the HLp-DNA complex. HLp alone and the 30-bp-GC40 DNA fragment eluted as single peaks, with molecular weights matching their theoretical values (Fig. 3D). Upon incubation of HLp with 30-bp-GC40 DNA, a new peak appeared, corresponding to a molecular weight of 69.2 ± 5.7 kDa, consistent with a complex consisting of an HLp tetramer bound to two 30-bp DNA duplexes.

### Crystal structures of HLp bound to DNA

To gain structural insight into the interaction between HLp and DNA, we co-crystallized HLp and the 30-bp-GC40 fragment. Diffraction data were collected for two different crystal forms, HLp-DNA_1 and HLp-DNA_2, processed to resolutions of 2.10 Å and

1.90 Å, respectively (Table S3). Both structures were solved by molecular replacement using the DNA-free HLp structure as the search model. In both HLp–DNA crystal forms, the DNA fragments appear to form continuous double helices winding throughout the crystal lattice, each revealing a distinct interface between HLp and the DNA (Fig. 4A and B). In both structures, the seemingly continuous DNA bases within central segments (16-bp dsDNA for HLp-DNA_1 and 15-nt ssDNA for HLp-DNA_2) were well resolved in the electron density maps, whereas the first six N-terminal residues of HLp were not visible.

**Fig. 4.**
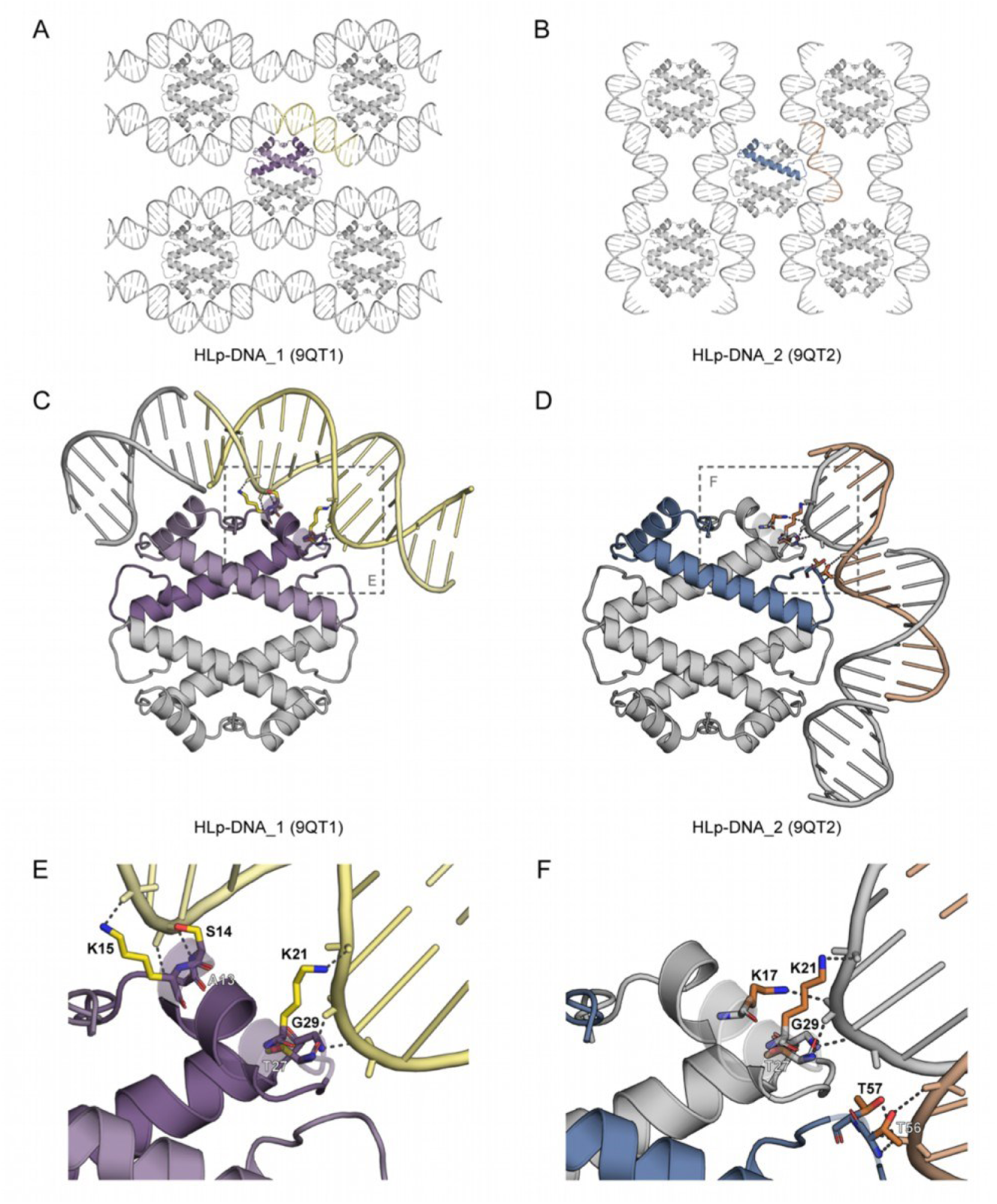
Crystal structures of DNA-bound HLp. **A,B.** Crystal packing of HLp-DNA_1 (**A**) and HLp-DNA_2 (**B**) showing selected symmetry mates within 20 Å. **C, D.** Crystal structures of HLp-DNA_1 (PDB: 9QT1) (**C**) and HLp-DNA_2 (PDB: 9QT2) (**D**) shown in cartoon representation. The framed regions in panels **C** and **D** correspond to the magnified views shown in **E** and **F**, respectively. **E, F.** Residues involved in DNA binding are shown as sticks, and protein-DNA interactions are depicted as dashed lines. In all panels, the contents of the asymmetric unit are shown in color, and selected symmetry mates in gray.

In HLp-DNA_1, the ASU contains an HLp dimer, which assembles into tetramers by crystallographic symmetry, and a 16-bp dsDNA segment of the seemingly infinite DNA helices winding throughout the crystal (Fig. 4A). As observed in other histone-DNA complexes, binding is mediated primarily through interactions between basic or polar side chains of HLp and the phosphate backbone of the DNA. On HLp, the interface spans the two monomers in the ASU, across the dimer. In each monomer, this interface involves residues A13, S14, and K15, which correspond to the “paired end of helices” motif in eukaryotic histones, as well as K21 in helix α1, and T27 and G29 in loop l1 (Fig. 4C and E)^58^.

In HLp-DNA_2, the ASU comprises a single HLp monomer, which forms tetramers via crystallographic symmetry, and a 15-nt ssDNA segment of the apparently endless DNA (Fig. 4B). Although the overall crystal packing is similar to HLp-DNA_1, it exhibits a complementary DNA binding mode, with the dsDNA engaged along the HLp tetramerization interface. Therein, in both HLp dimers, the monomers bind to DNA via two sets of interactions. The first corresponds to the “β-bridge” motif in eukaryotic histones and involves T27 and G29 in loop l1, as well as T56 and T57 in loop l2, while the second set involves K17 and K21 from helix α1 (Fig. 4D and F)^58^.

The two DNA-bound crystal structures thereby reveal complementary, but overlapping DNA-binding interfaces distributed along the orbicular surface of the HLp tetramer, corresponding to the canonical interaction motifs of archaeal and eukaryotic histones^16, 58^. Taken together, these findings suggest that HLp is able to bind DNA through a wrapping mechanism.

### Molecular dynamics (MD) simulation

The two crystal structures suggested a bridging mode of DNA binding by HLp. However, given the distinct yet overlapping interfaces observed, we hypothesized that HLp might also support a wrapping mode, in which the tetramer could engage a DNA segment across the entire DNA-binding interface. To investigate whether HLp indeed exhibits both bridging and wrapping modes, and to explore their relative stability and potential interconversion, we performed all-atom MD simulations. Using the HLp- DNA_1 and HLp-DNA_2 structures as templates, we generated two starting models: a wrapping model, consisting of an HLp tetramer wrapped by a 67-bp dsDNA fragment, and a bridging model, in which an HLp tetramer engages two separate 32-bp dsDNA fragments (Fig. 5A and Fig. S6). For each model, we performed two independent 1 µs all-atom simulations, which yielded highly consistent trajectories (SI movie 1 and 2). For subsequent analyses, one representative trajectory per model was examined in detail.

**Fig. 5.**
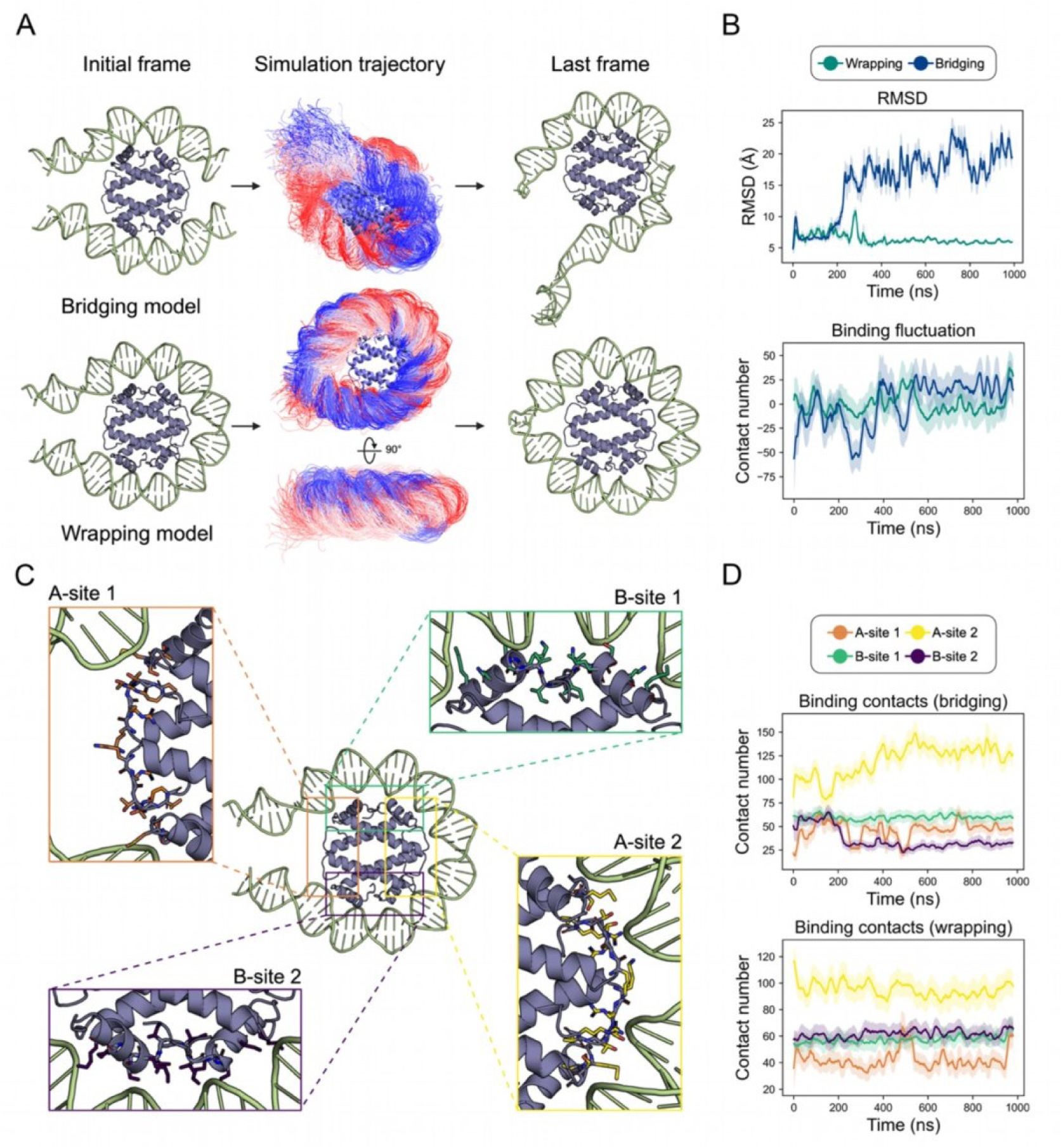
Molecular dynamics simulations of the wrapping and bridging modes. **A.** Visualization of the starting model (left), the conformational ensemble derived from the simulation (middle), and the structure of the last frame (right). **B.** Overall characterization of the simulations of the two proposed binding modes, showing the RMSD of the DNA backbones (top) and binding fluctuations (bottom). Binding fluctuation is defined as the deviation of the contact number in each frame from the average contact number across all frames. Each plot is smoothed with a window size of 100, with green and blue lines represent the wrapping and bridging models, respectively. **C.** Structures of the four defined binding sites: A-site 1 (orange), A-site 2 (yellow), B-site 1 (green), and B-site 2 (purple). Residues involved in each site are shown as sticks and listed in Table S4. **D.** Binding contacts for the four defined binding sites throughout the simulation for the wrapping (top) and bridging (bottom) models. Line colors correspond to those in panel C. Binding contacts are defined as any pair of heavy atoms within 4 Å.

The wrapping model showed high stability, with a DNA backbone RMSD maintained at approximately 6 Å throughout the simulation (Fig. 5B). The final frame of the simulation shows the HLp tetramer fully wrapped by ∼60 bp of dsDNA (Fig. 5A). In contrast, the bridging model exhibited substantial conformational rearrangements, characterized by a progressive increase in protein-DNA contacts and a spontaneous shift toward the wrapping configuration (Fig. 5A and B). Consistently, the DNA potential energy of the wrapping model is lower than the bridging model (Fig. S7A), suggesting that the wrapping interaction provides greater thermodynamic stability through an expanded protein-DNA interface.

To quantify HLp-DNA contacts, we calculated atom-atom interactions between protein and DNA heavy atoms for both the bridging and wrapping models (Fig. S8 and S9). Most contacts involved the DNA phosphate backbone and sugar moieties (Fig. S7B), consistent with the non-sequence-specific binding observed in crystal structures. HLp side chains contributed slightly more to DNA interactions than backbone atoms (Fig. S6C). Across both models, residues located in α1 and loops l1 and l2 exhibited an average of more than three DNA contacts (Fig. S8 and S9). In particular, K54 and R59 in loop l2 formed persistent hydrogen bonds with the DNA throughout the simulations (Fig. S10).

To dissect the spatial dynamics of binding, we defined four major DNA-binding regions on the HLp tetramer: A-sites 1 and 2, each comprising two “β-bridge” motifs with their flanking DNA-binding residues; and B-sites 1 and 2, each centered on a “paired end of helices” motif along with adjacent DNA-interacting residues (Fig. 5C and Table S4). In the bridging model, DNA remained stably associated with B-sites 1 and 2, but progressively engaged A-site 2, and to a lesser extent A-site 1, as it transitioned into the wrapping mode (Fig. 5D and Fig. S11). In the wrapping model, local interactions at all four sites were stably maintained or even enhanced over time (Fig. 5D and Fig. S12). Notably, fluctuations in contact number at A-site 1 in both models appear to result from steric interference at the DNA termini.

To probe the unwrapping dynamics of the HLp-DNA complex, we performed steered molecular dynamics (SMD) simulations (Fig. S13). Unwrapping occurred in two distinct phases: initial disruption at B-site 2, followed by disengagement at A-site 2, yielding a partially unwrapped intermediate ensemble. The unwrapping energy landscape revealed two main barriers: ∼500 kJ/mol at B-site 2 and ∼400 kJ/mol at A-site 2. Energy decomposition analysis identified five residues — R59, K21, K15, S14, and K17 — as major contributors to DNA binding, collectively accounting for 52% of the total binding energy. These findings are consistent with the hydrogen bond and contact analyses from the unbiased simulations and highlight the cooperative, multivalent nature of HLp-DNA interactions. The high energetic cost of unwrapping suggests that dissociation is a regulated, stepwise process, reinforcing the stability of the wrapped state.

Together, our simulations support both wrapping and bridging as viable DNA-binding modes for HLp. However, the wrapping mode is energetically favored and yields more extensive, stable interactions — pointing to its likely predominance under physiological conditions.

### HLp is wrapped by DNA *in vitro*

To investigate the mechanism of DNA binding by HLp *in vitro*, we employed a series of well-established biochemical and biophysical assays. We first used a MNase digestion assay, which measures the extent to which protein-bound DNA is protected from nucleolytic cleavage. The 600-bp dsDNA fragment was incubated with HLp and subsequently digested with increasing concentrations of MNase. The resulting digestion pattern revealed a characteristic ladder of fragments ranging from 35 to 72 bp (Fig. 6A), indicating that HLp protects bound DNA from enzymatic degradation. This protective effect is reminiscent of that observed for the nucleosomal archaeal histone HMfB, which served as a positive control. However, the HLp-protected fragments differ in both size and regularity: while HMfB consistently generates ∼30-bp increments, the fragment sizes protected by HLp are more heterogeneous. These observations suggest that although both proteins compact DNA, they do so through distinct binding modes.

**Fig. 6.**
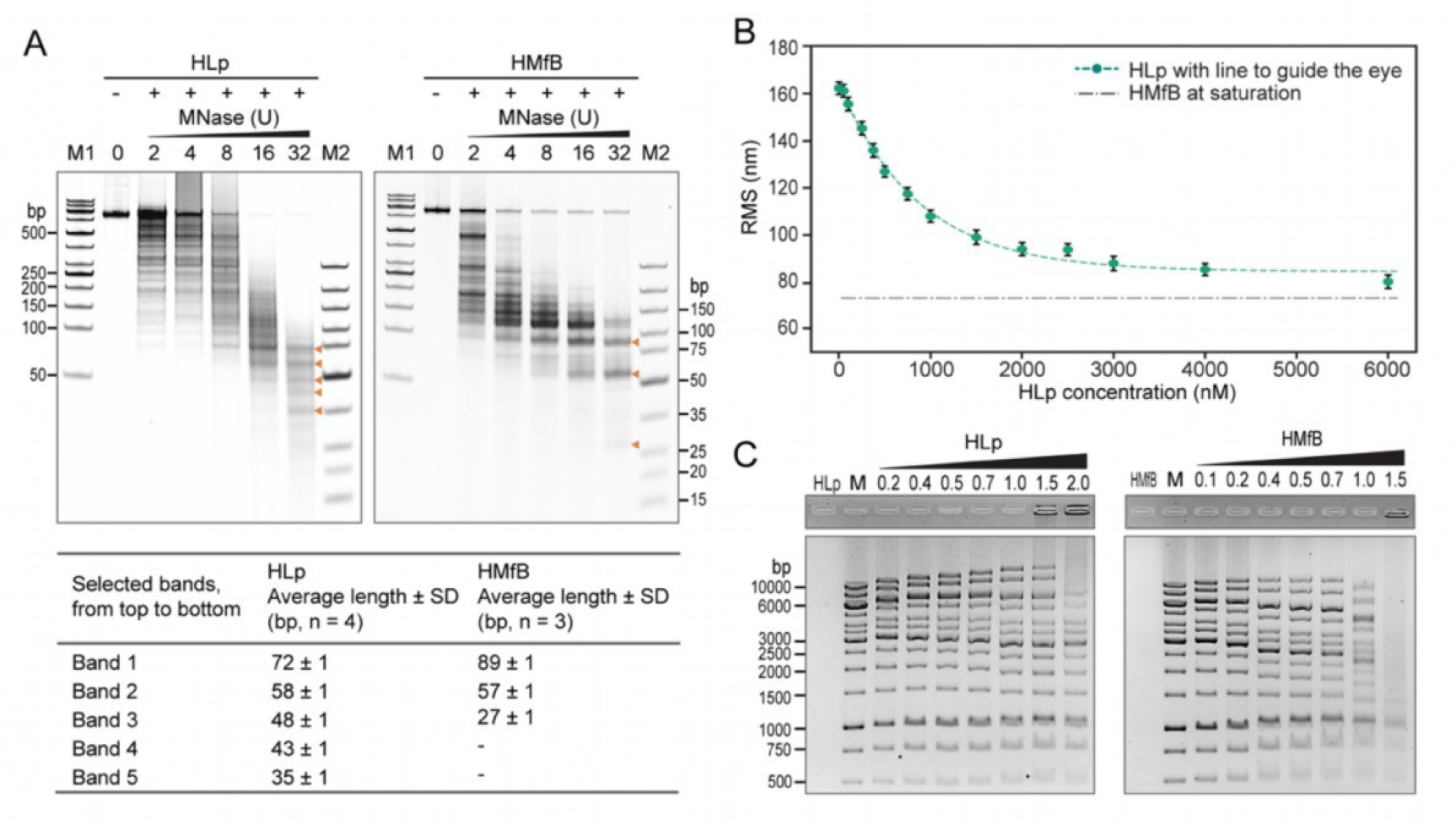
DNA wraps around HLp. **A.** MNase digestion assay analyzing the protection of the 600-bp-GC40 DNA fragment by HLp in comparison to HMfB. Increasing MNase concentrations used in the assay are indicated. The lengths of DNA fragments obtained at the highest MNase concentration were determined by densitometric analysis, using the molecular weight marker M2 (GeneRuler Ultra Low Range DNA Ladder, ThermoFisher Scientific) as the standard. **B.** TPM experiment with HLp and the 685-bp DNA. Each measurement point represents the average of triplicate measurements, with error bars indicating standard deviations. The line connecting the points was generated by fitting the means to a logistic function. **C.** EMSA analyzing the binding of HLp and HMfB (control) to the DNA fragments of the GeneRuler 1 kb Ladder (ThermoFisher Scientific). Increasing protein concentrations (lanes 3-9), indicated as protein-to-DNA mass ratios, were incubated with the corresponding DNA and analyzed on a 1% agarose gel.

To directly monitor HLp-induced DNA compaction, we performed TPM experiments using the 685-bp linear DNA fragment. TPM enables real-time monitoring of protein-DNA interactions by measuring the Brownian motion of a DNA-tethered polystyrene bead; changes in DNA flexibility or contour length, such as those caused by protein-induced compaction, result in measurable shifts in root mean square (RMS) displacement of the bead. Upon addition of HLp, we observed a progressive, concentration-dependent decrease in RMS values, indicating compaction of the DNA (Fig. 6B). Saturation was reached at ∼6000 nM HLp with a final RMS value of ∼80 nm, closely matching the values observed for the archaeal histones HMfA and HMfB, suggesting that HLp wraps DNA rather than bridging it^59^.

To investigate whether HLp induces DNA condensation across a range of DNA sizes, we performed ladder EMSA assays with linear dsDNA fragments of increasing length. HLp altered the electrophoretic mobility of DNA on agarose gels in a size-dependent manner, with fragments longer than 2000 bp migrating faster, while those shorter than 1000 bp showed reduced mobility (Fig. 6C). These results are consistent with protein-induced changes in DNA conformation and are qualitatively similar to those observed with HMfB, although HMfB required lower protein concentrations and produced more pronounced mobility shifts^17^.

We next examined whether HLp binding affects DNA topology using a topoisomerase I relaxation assay. In the presence of HLp, the relaxed plasmid DNA became progressively supercoiled in a protein concentration-dependent manner, indicating the introduction of topological strain upon HLp binding (Fig. S14A). This behavior mirrors that of HMfB, albeit with a less pronounced effect.

Finally, we tested whether HLp promotes DNA end-joining using a ligase-mediated circularization assay. In this assay, short DNA fragments are circularized by T4 DNA ligase and non-circularized linear species are digested by T5 exonuclease. While HMfB efficiently promotes the circularization of DNA monomers due to its pronounced DNA-bending activity, HLp had only modest effects, even at high concentrations (Fig. S14B). In contrast, HLp favored the formation of linear DNA multimers as protein concentration increased, indicative of open-ended protein-DNA complexes formed upon binding (Fig. S14B).

Taken together with the crystal structures and molecular dynamics simulations, our *in vitro* data support a model in which HLp wraps and compacts DNA through a mechanism that is distinct from the bending mode employed by the bacterial histone HBb and from the nucleosome assembly of eukaryotic histones, differing in both its dynamics and topological effects.

### HLp binds to genomic DNA *in vivo*

Due to the lack of facilities for handling *Leptospira* strains, we used *E. coli* as a heterologous model to investigate the effects of HLp on genomic DNA *in vivo* using light microscopy. HLp was expressed in *E. coli* BL21(DE3), while the archaeal histone HMfB was expressed in *E. coli* Mutant56(DE3) as a positive control. Ubiquitin from *Caldiarchaeum subterraneum* (*Cs*Ub), which lacks DNA-binding activity, served as a negative control^32^. Because expression levels varied significantly between constructs, the strain showing the highest expression level for each protein was selected for further analysis.

Growth curves were recorded to determine the onset of the stationary phase and to assess potential effects of protein expression on cell proliferation (Fig. S15). HLp and *Cs*Ub expression in *E. coli* BL21(DE3) led to a modest increase in doubling time and a reduction in final cell density compared to uninduced controls. However, similar effects were observed in cells carrying empty vectors, suggesting that these changes were not specific to HLp or *Cs*Ub overexpression. In contrast, HMfB expression had no measurable impact on cell growth. To minimize variability due to nucleoid dynamics during the cell cycle, microscopy samples were collected in early stationary phase, when cells had exited active division.

Microscopic analysis revealed that HLp expression markedly alters nucleoid organization. DAPI staining of HLp-expressing cells demonstrated clear elongation and a substantial increase in nucleoid volume relative to controls (Fig. 7), consistent with impaired chromosome condensation and/or interference with cell division. These effects were already evident in the absence of IPTG, likely due to basal expression from the leaky T7 promoter. Upon overexpression of HLp, the cytoplasm of HLp- expressing cells was predominantly occupied by decondensed genomic DNA, a phenotype also observed upon HMfB overexpression, suggesting that HLp disrupts the chromatin organization of the *E. coli* cells through its DNA-binding ability. Segmentation of the DAPI fluorescence signals in HMfB-expressing cells exhibited single, well-defined regions with clear contours. In contrast, segmentation of HLp- expression cells resulted in fluorescence signals dispersed across multiple regions with diffuse contours, indicating a more heterogeneous distribution of these nucleoid regions. Combined with TPM assay results, which showed HLp-induced DNA compaction only at high protein concentrations, these findings suggest that HLp induces localized DNA condensation and reveal subtle differences in chromatin organization between the two histones.

**Fig. 7.**
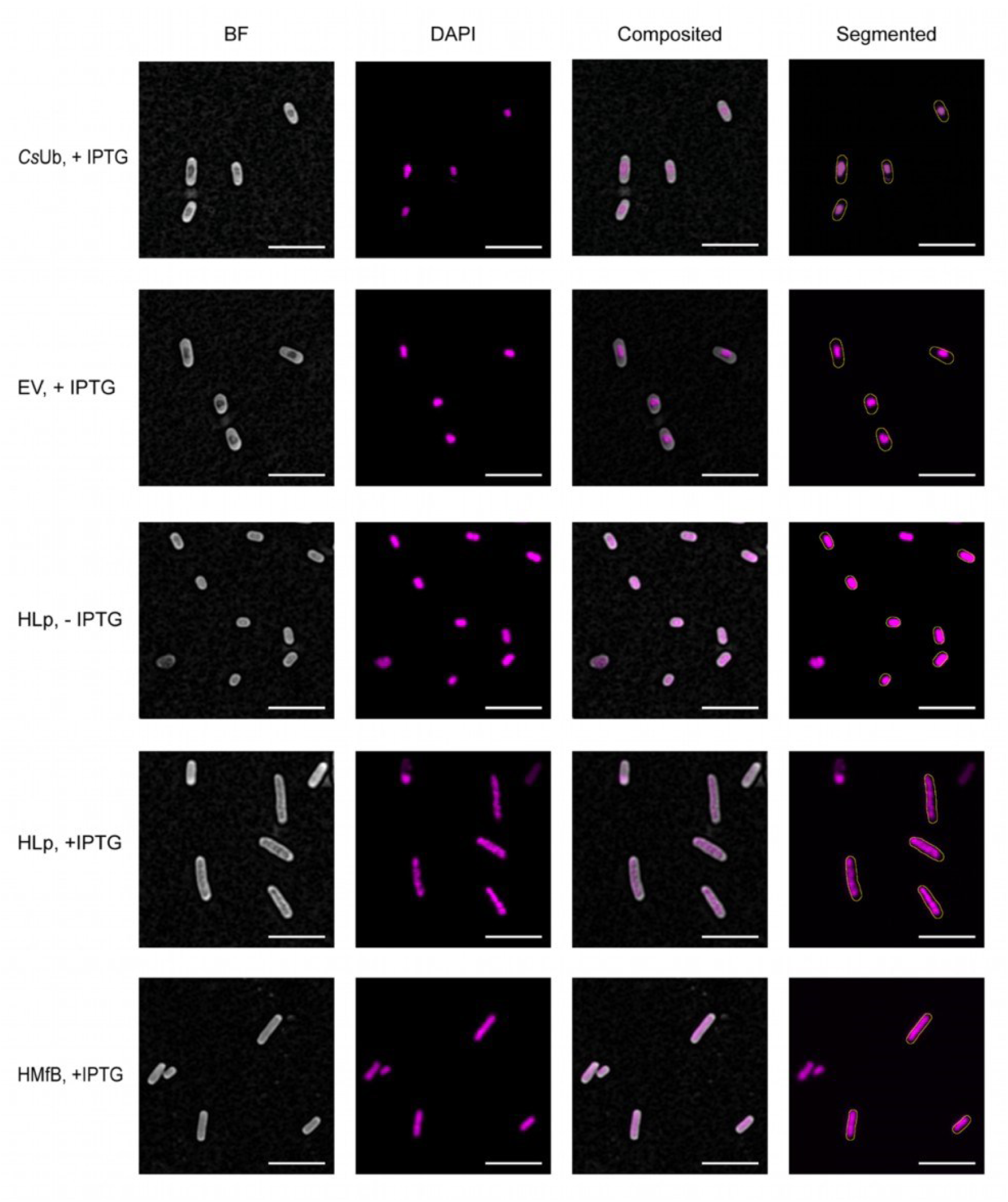
**HLp binds to genomic DNA *in vivo*.** Airyscan imaging of *E. coli* cells expressing HLp and HMfB (positive control) with DAPI dye, compared to negative controls transformed with the empty vector (EV), the vector expressing *Cs*Ub, and the uninduced HLp sample (-IPTG). The left column presents the brightfield image with subtracted background. The second column displays the maximum intensity projection of fluorescence. The third one is the overlap, and the last column shows the segmented bacteria outline on top of the fluorescence channel. Scale bar: 5 µm.

## Discussion

In this study, we present a comprehensive analysis of the bacterial histone HLp from *L. perolatii*, integrating structural, biophysical, and functional data to elucidate its role in DNA binding and compaction. Our findings reveal that HLp assembles into a unique, DNA-wrapped tetramer, expanding the known repertoire of bacterial histone architectures, and providing new insights into prokaryotic chromatin organization. This represents the first experimental characterization of a bacterial histone belonging to the FtF subgroup of α3 histones, previously identified bioinformatically as one of the largest subfamilies of prokaryotic histones^9^.

Structurally, HLp closely resembles the archaeal FtF histone HTkC from *T. kodakarensis*, consistent with bioinformatic predictions based on CLANS clustering, where the two histones are positioned adjacent to each other. Both proteins adopt a characteristic histone fold comprising three α-helices connected by two loops, yet differ notably from nucleosomal histones in their truncated α2 and α3 helices. This truncation is also found in HBb, the only other experimentally studied bacterial histone, highlighting a common structural adaptation within α3 histones^20, 24^. Unlike HBb, which strictly forms dimers, HLp and HTkC assemble into tetramers even in the absence of DNA, mediated by highly conserved residues situated at the C-terminal region of α2, loop l2, and helix α3. This unique oligomerization interface appears to be a defining feature of FtF histones and sets them apart from other prokaryotic histone families, including archaeal nucleosomal histones, which typically assemble into higher-order oligomers only upon DNA binding. Given their structural similarity and conserved tetramerization interface, we anticipate that HTkC may also wrap DNA similarly to HLp.

The HLp tetramer displays a continuous band of positive surface charge encircling the complex, consistent with its role in non-specific DNA binding. EMSA and MST assays confirm that HLp binds dsDNA without sequence specificity, with strongest affinity for DNA of ∼40% GC content, which is similar to the native genomic composition of *L. perolatii*. Crystal structures of HLp-DNA complexes reveal the HLp tetramer to be decorated with distinct DNA-binding interfaces along its perimeter, including motifs corresponding to the canonical “paired end of helices” and “β-bridge” motifs^58^. Thereby, their spatial arrangement in HLp resembles that of eukaryotic and archaeal histones.

While each of the two individual DNA-bound crystal structures might hint at a DNA-bridging mode, their structural superposition and the location of the individual DNA-binding motifs suggested a potential DNA-wrapping mode for HLp. Of these two modes, MD simulations favor the wrapping mode as the energetically preferred configuration. Even simulations initiated in the bridging conformation exhibited spontaneous progression toward wrapping. Notably, DNA wrapping involves ∼60 bp encircling the tetramer, consistent with structural predictions. However, we cannot rule out that HLp also bridges DNA *in vivo*, potentially in concert with other factors.

Biochemical assays further support the wrapping-based binding mode. In MNase digestion assays, HLp protects DNA from nucleolytic cleavage, producing irregularly spaced fragments (35-72 bp) distinct from the regular 30-bp pattern generated by HMfB and the ∼147-bp protection seen in eukaryotic nucleosomes. These fragment sizes correspond approximately to half and full turns of DNA around the HLp tetramer, indicating protection via wrapping but with flexible nucleosome-like spacing. Additional *in vitro* assays, including TPM, topoisomerase relaxation and ladder EMSA, demonstrate that HLp compacts DNA and alters its topology, albeit less efficiently than HMfB. HLp does not promote the formation of circular DNA monomers in the circularization assay, suggesting that it does not bring DNA ends into close proximity for ring closure under the tested conditions.

Expression of HLp in *E. coli* leads to dramatic reorganization of the nucleoid. DAPI staining reveals marked nucleoid decondensation and increased cellular length, indicative of disrupted DNA compaction and/or interference with cell division. These phenotypes closely mirror those observed upon HMfB expression, although the nucleoid appears more homogeneous with HMfB than with HLp, suggesting differences in chromatin architecture. These *in vivo* findings support the hypothesis that HLp functions as a global chromatin organizer, similar to its homolog in *L. interrogans*, where the FtF histone is among the most highly expressed and essential proteins^20^. Notably, neither *L. interrogans* nor *L. perolatii* encodes other histone homologs or canonical bacterial NAPs, further indicating that HLp serves as the principal chromatin component in these species. In contrast, species such as *B. bacteriovorus* encode two distinct histones (HBb and Bd3044) as well as NAPs such as HU, underscoring lineage-specific diversity in bacterial chromatin organization strategies^20, 24^. Consistent with HLp’s proposed central role, genomic analysis reveals that in *L. perolatii*, the *hlp* gene resides within a conserved locus adjacent to genes encoding SMC, MAP, and a putative transmembrane protein (Fig. S1B). This synteny is preserved in *L. interrogans* and other *Leptospira* species, suggesting a functionally co-evolved genomic module (Fig. S1B). Such an arrangement raises the possibility that HLp cooperates closely with SMC to maintain proper chromosome structure and segregation, while the neighboring MAP enzyme, responsible for cleaving the initiating methionine from nascent proteins, may support efficient maturation of HLp, SMC, and other critical factors involved in nucleoid dynamics.

From an evolutionary perspective, HLp is only the second bacterial histone characterized structurally and functionally, after the dimeric histone HBb. While bacterial dimeric histones like HBb appear restricted to bacteria and nucleosomal histones are found exclusively in archaea and eukaryotes, FtF histones uniquely span both archaeal and bacterial domains. Our bioinformatic analysis demonstrates that bacterial and archaeal FtF histones form distinct and clearly separated clusters, supporting the hypothesis that FtF histones may trace their ancestry back to the last universal common ancestor (LUCA). Alternatively, the broad distribution of FtF histones across archaeal phyla, including Asgard archaea, and their sparse, patchy occurrence in bacteria, suggest a scenario involving multiple independent horizontal gene transfer events from archaea into specific bacterial lineages^9^. Thus, the evolutionary history of FtF histones likely reflects a combination of ancestral origin, lineage-specific adaptations, and recurrent gene-transfer events.

In summary, our work expands the known diversity of bacterial histones and establishes HLp as a DNA-wrapping histone that assembles into a stable homotetramer. Rather than being simplified ancestors of eukaryotic histones, prokaryotic histones represent structurally diverse, functionally adaptable proteins that contribute to genome organization through distinct DNA-binding modes. Beyond structural and mechanistic insight, future studies will need to explore how these histones function in their native cellular contexts — how they interact with other histone variants, cooperate with or substitute for NAPs, and participate in processes such as transcription, replication, and genome repair.

## Data availability

Coordinates and structure factors of the crystal structures have been deposited in the PDB under entry numbers 9QT0 (DNA-free HLp), 9QT1 (HLp-DNA_1) and 9QT2 (HLp- DNA_2).

For the MD analysis, all configuration files, trajectory files, movies, and analysis scripts are deposited at Zenodo (http://zenodo.org/) and accessible with the DOI 10.5281/zenodo.15188379.

The TPM data is deposited in the 4TU repository (https://data.4tu.nl) and accessible with the DOI 10.4121/5b604dd5-5498-46aa-b437-775e957f93e3.

## Acknowledgements

We thank the staff of Beamline X10SA of the Swiss Light Source (PSI, Villigen, Switzerland) for excellent technical support. We are grateful to Reinhard Albrecht for assistance with crystallization and crystallographic data collection. We extend our thanks to Linxuan Li (Dept. of Integrative Evolutionary Biology, MPI for Biology Tübingen, Germany) and Agnes Henschen (Dept. of Algal Development and Evolution, MPI for Biology Tübingen, Germany) for their support with light microscopy sample preparation. We thank Pedro Escudeiro (Dept. of Protein Evolution, MPI for Biology Tübingen, Germany) for proofreading the manuscript. We acknowledge the HPC system Raven at the Max Planck Computing and Data Facility for the performed computational work.

## Author Contributions

Conceived and designed the experiments: A.N.L., B.H.A., V.A., Y.H.

Performed the experiments: A.P., H.R., K.B., S.S., Y.H.

Performed MD simulations: K.Q., Y.Z.

Performed bioinformatic analyses: V.A.

Analyzed the data: A.P., B.H.A., M.D.H., S.S., R.T.D, V.A., Y.H.

Wrote the paper: B.H.A., V.A., Y.H., with contributions from the other authors.

## References

1. Arents G, Burlingame RW, Wang BC, Love WE, Moudrianakis EN. The Nucleosomal Core Histone Octamer at 3.1-a Resolution - a Tripartite Protein Assembly and a Left-Handed Superhelix. P Natl Acad Sci USA 88, 10148–10152 (1991).

2. Luger K, Mader AW, Richmond RK, Sargent DF, Richmond TJ. Crystal structure of the nucleosome core particle at 2.8 angstrom resolution. Nature 389, 251–260 (1997).

3. Kornberg RD, Thomas JO. Chromatin structure; oligomers of the histones. Science 184, 865–868 (1974).

4. Allan J, Hartman PG, Crane-Robinson C, Aviles FX. The structure of histone H1 and its location in chromatin. Nature 288, 675–679 (1980).

5. Li W, et al. Structural basis for linker histone H5–nucleosome binding and chromatin fiber compaction. Cell Research 34, 707–724 (2024).

6. Luger K, Richmond TJ. The histone tails of the nucleosome. Curr Opin Genet Dev 8, 140–146 (1998).

7. Jenuwein T, Allis CD. Translating the histone code. Science 293, 1074–1080 (2001).

8. Hocher A, Warnecke T. Nucleosomes at the Dawn of Eukaryotes. Genome Biol Evol 16, evae029 (2024).

9. Schwab S, et al. Histones and histone variant families in prokaryotes. Nature Communications 15, 7950 (2024).

10. Henneman B, van Emmerik C, van Ingen H, Dame RT. Structure and function of archaeal histones. PLoS Genet 14, e1007582 (2018).

11. Stevens KM, et al. Histone variants in archaea and the evolution of combinatorial chromatin complexity. P Natl Acad Sci USA 117, 33384–33395 (2020).

12. Dame RT. The role of nucleoid-associated proteins in the organization and compaction of bacterial chromatin. Mol Microbiol 56, 858–870 (2005).

13. Dorman CJ, Deighan P. Regulation of gene expression by histone-like proteins in bacteria. Current Opinion in Genetics & Development 13, 179–184 (2003).

14. Dillon SC, Dorman CJ. Bacterial nucleoid-associated proteins, nucleoid structure and gene expression. Nature Reviews Microbiology 8, 185–195 (2010).

15. Dame RT. Special Issue: Role of Bacterial Chromatin in Environmental Sensing, Adaptation and Evolution. Microorganisms 9, (2021).

16. Mattiroli F, et al. Structure of histone-based chromatin in Archaea. Science 357, 609–612 (2017).

17. Sandman K, Krzycki JA, Dobrinski B, Lurz R, Reeve JN. HMf, a DNA-binding protein isolated from the hyperthermophilic archaeon Methanothermus fervidus, is most closely related to histones. Proc Natl Acad Sci U S A 87, 5788–5791 (1990).

18. Sanders TJ, et al. Extended Archaeal Histone-Based Chromatin Structure Regulates Global Gene Expression in Thermococcus kodakarensis. Front Microbiol 12, 681150 (2021).

19. Ofer S, et al. DNA-bridging by an archaeal histone variant via a unique tetramerisation interface. Communications Biology 6, 968 (2023).

20. Hocher A, et al. Histones with an unconventional DNA-binding mode in vitro are major chromatin constituents in the bacterium Bdellovibrio bacteriovorus. Nat Microbiol 8, 2006–2019 (2023).

21. Alva V, Lupas AN. Histones predate the split between bacteria and archaea. Bioinformatics 35, 2349–2353 (2019).

22. Qiu Y, et al. The crystal structure of Aq_328 from the hyperthermophilic bacteria Aquifex aeolicus shows an ancestral histone fold. Proteins 62, 8–16 (2006).

23. Paratsaphan S, et al. Characterization of a novel peptide from pathogenic leptospira and its cytotoxic effect. Pathogens 9, 906 (2020).

24. Hu Y, et al. Bacterial histone HBb from Bdellovibrio bacteriovorus compacts DNA by bending. Nucleic Acids Res 52, 8193–8204 (2024).

25. Camacho C, et al. BLAST+: architecture and applications. BMC bioinformatics 10, 1–9 (2009).

26. Sayers EW, et al. Database resources of the National Center for Biotechnology Information in 2025. Nucleic acids research 53, D20–D29 (2025).

27. Steinegger M, Söding J. MMseqs2 enables sensitive protein sequence searching for the analysis of massive data sets. Nature biotechnology 35, 1026–1028 (2017).

28. Frickey T, Lupas A. CLANS: a Java application for visualizing protein families based on pairwise similarity. Bioinformatics 20, 3702–3704 (2004).

29. Zimmermann L, et al. A completely reimplemented MPI bioinformatics toolkit with a new HHpred server at its core. Journal of molecular biology 430, 2237–2243 (2018).

30. Oberg N, Zallot R, Gerlt JA. EFI-EST, EFI-GNT, and EFI-CGFP: enzyme function initiative (EFI) web resource for genomic enzymology tools. Journal of molecular biology 435, 168018 (2023).

31. Jumper J, et al. Highly accurate protein structure prediction with AlphaFold. Nature 596, 583–589 (2021).

32. Fuchs ACD, Maldoner L, Wojtynek M, Hartmann MD, Martin J. Rpn11-mediated ubiquitin processing in an ancestral archaeal ubiquitination system. Nature Communications 9, 2696 (2018).

33. Bogomolovas J, Simon B, Sattler M, Stier G. Screening of fusion partners for high yield expression and purification of bioactive viscotoxins. Protein Expr Purif 64, 16–23 (2009).

34. Kabsch W. Xds. Acta Crystallogr D Biol Crystallogr 66, 125–132 (2010).

35. 35. Tickle I, et al. Staraniso. Cambridge, United Kingdom: Global Phasing Ltd 923, (2018).

36. Evans R, et al. Protein complex prediction with AlphaFold-Multimer. biorxiv, 2021.2010. 2004.463034 (2021).

37. Vagin A, Teplyakov A. MOLREP: an automated program for molecular replacement. J Appl Crystallogr 30, 1022–1025 (1997).

38. Emsley P, Cowtan K. Coot: model-building tools for molecular graphics. Acta Crystallogr D Biol Crystallogr 60, 2126–2132 (2004).

39. Murshudov GN, et al. REFMAC5 for the refinement of macromolecular crystal structures. Acta Crystallogr D Biol Crystallogr 67, 355–367 (2011).

40. Van Der Spoel D, Lindahl E, Hess B, Groenhof G, Mark AE, Berendsen HJ. GROMACS: fast, flexible, and free. Journal of computational chemistry 26, 1701–1718 (2005).

41. Huang J, et al. CHARMM36m: an improved force field for folded and intrinsically disordered proteins. Nature methods 14, 71–73 (2017).

42. Mark P, Nilsson L. Structure and dynamics of the TIP3P, SPC, and SPC/E water models at 298 K. The Journal of Physical Chemistry A 105, 9954–9960 (2001).

43. Parrinello M, Rahman A. Polymorphic transitions in single crystals: A new molecular dynamics method. Journal of Applied physics 52, 7182–7190 (1981).

44. Bussi G, Donadio D, Parrinello M. Canonical sampling through velocity rescaling. The Journal of chemical physics 126, (2007).

45. Darden T, York D, Pedersen L. Particle mesh Ewald: An N log (N) method for Ewald sums in large systems. Journal of chemical physics 98, 10089–10089 (1993).

46. Hess B, Bekker H, Berendsen HJ, Fraaije JG. LINCS: A linear constraint solver for molecular simulations. Journal of computational chemistry 18, 1463–1472 (1997).

47. Park S, Schulten K. Calculating potentials of mean force from steered molecular dynamics simulations. The Journal of chemical physics 120, 5946–5961 (2004).

48. Humphrey W, Dalke A, Schulten K. VMD: visual molecular dynamics. Journal of molecular graphics 14, 33–38 (1996).

49. Michaud-Agrawal N, Denning EJ, Woolf TB, Beckstein O. MDAnalysis: a toolkit for the analysis of molecular dynamics simulations. Journal of computational chemistry 32, 2319–2327 (2011).

50. Schindelin J, et al. Fiji: an open-source platform for biological-image analysis. Nature methods 9, 676–682 (2012).

51. Henneman B, Heinsman J, Battjes J, Dame RT. Quantitation of DNA-Binding Affinity Using Tethered Particle Motion. Methods Mol Biol 1837, 257–275 (2018).

52. 52. Huff J. The Airyscan detector from ZEISS: confocal imaging with improved signal-to-noise ratio and super-resolution. (eds). Nature Publishing Group US New York (2015).

53. Thibeaux R, et al. Deciphering the unexplored Leptospira diversity from soils uncovers genomic evolution to virulence. Microbial genomics 4, e000144 (2018).

54. Shine J, Dalgarno L. The 3′-terminal sequence of Escherichia coli 16S ribosomal RNA: complementarity to nonsense triplets and ribosome binding sites. Proceedings of the National Academy of Sciences 71, 1342–1346 (1974).

55. Decanniere K, Babu AM, Sandman K, Reeve JN, Heinemann U. Crystal structures of recombinant histones HMfA and HMfB from the hyperthermophilic archaeon Methanothermus fervidus. J Mol Biol 303, 35–47 (2000).

56. Baker NA, Sept D, Joseph S, Holst MJ, McCammon JA. Electrostatics of nanosystems: application to microtubules and the ribosome. Proceedings of the National Academy of Sciences 98, 10037–10041 (2001).

56. Bailey KA, Pereira SL, Widom J, Reeve JN. Archaeal histone selection of nucleosome positioning sequences and the procaryotic origin of histone-dependent genome evolution. J Mol Biol 303, 25–34 (2000).

58. Arents G, Moudrianakis EN. Topography of the histone octamer surface: repeating structural motifs utilized in the docking of nucleosomal DNA. Proc Natl Acad Sci U S A 90, 10489–10493 (1993).

59. Henneman B, et al. Mechanical and structural properties of archaeal hypernucleosomes. Nucleic Acids Res 49, 4338–4349 (2021).

