## Supplemental information for "DNA Wrapping by a Tetrameric Bacterial Histone"

[#] Joint second authors

*To whom correspondence should be addressed

### Supplementary Tables

#### Supplementary Table S1
#### Oligonucleotides used in this work.

| Oligo name | Sequence (5' to 3') | Application |
|---|---|---|
| 30-bp-GC30-F | TTTAAAACGCTTTAAAACGCTTTAAAACGC | EMSA |
| 30-bp-GC30-R | GCGTTTTAAAGCGTTTTAAAGCGTTTTAAA | EMSA |
| 30-bp-GC40-F | TTTAAAGCCGTTTAAAGCCGTTTAAAGCCG | SEC-MALS, EMSA, Crystallization |
| 30-bp-GC40-R | CGGCTTTAAACGGCTTTAAACGGCTTTAAA | SEC-MALS, EMSA, Crystallization |
| 30-bp-GC50-F | TTAAAGCCCGTTAAAGCCCGTTAAAGCCCG | EMSA |
| 30-bp-GC50-R | CGGGCTTTAACGGGCTTTAACGGGCTTTAA | EMSA |
| 30-bp-GC60-F | TTAAGCCCCGTTAAGCCCCGTTAAGCCCCG | EMSA |
| 30-bp-GC60-R | CGGGGCTTAACGGGGCTTAACGGGGCTTAA | EMSA |
| 80-bp-DNA-F | CCGTACTGTCGTCTGCGGCCTTTGATTATCA ATTAAAGCGTTCTACGGCGTTTTTGATCGCT CAACGTGCGGAGCTAGAT | EMSA |
| 80-bp-DNA-F[Cy5] | [Cyanine5]CCGTACTGTCGTCTGCGGCCTTTG ATTATCAATTAAAGCGTTCTACGGCGTTTTTG ATCGCTCAACGTGCGGAGCTAGAT | MST |
| 80-bp-DNA-R | ATCTAGCTCCGCACGTTGAGCGATCAAAAAC GCCGTAGAACGCTTTAATTGATAATCAAAGG CCGCAGACGACAGTACGG | EMSA, MST |
| pET-600-bp-GC40-F | CGCGAATTTTAACAAAATATTAACGTTTACA | MNase digestion |
| pET-600-bp-GC40-R | ATTCAGGTGAAAATATTGTTGATGCG | MNase digestion |
| 80-bp-GC40-F[Cy5] | [Cyanine5]TTTAAAGCCGTTTAAAGCCGTTTAA AGCCGTTTAAAGCCGTTTAAAGCCGTTTAAA GCCGTTTAAAGCCGTTTAAAGCCG | MST |
| 80-bp-GC40-R | CGGCTTTAAACGGCTTTAAACGGCTTTAAAC GGCTTTAAACGGCTTTAAACGGCTTTAAACG GCTTTAAACGGCTTTAAA | MST |
| 685-bp-DNA-F[Biotin] | [Biotin]TTACTTTCACCAGCGTTTCTGGGTGAG CAAAAACAG | TPM |

| | | |
|---|---|---|
| 685-bp-DNA-R[DIG] | [DIG]CCAAGTAGCGAAGCGAGCAGGACTGGGCGG | TPM |
| pET-240-bp-GC40-Fp | [Phos]TGCAATTTATTCATATCAGGATTATCA | Ligase-mediated circularization assay |
| pET-240-bp-GC40-Rp | [Phos]GCATAAACTTTTGCCATTCTCACC | Ligase-mediated circularization assay |

**Supplementary Table S2**
**Binding affinities ($K_d$) of HLp and HMfB to DNA substrates measured with MST.**

| Protein | DNA substrate | $K_d$ ± SD* (μM) |
|---|---|---|
| HLp | 80-bp DNA | 3.18 ± 1.06 |
| | 80-bp-GC40 DNA | 1.24 ± 0.14 |
| HMfB | 80-bp DNA | 0.58 ± 0.13 |
| | 80-bp-GC40 DNA | 0.31 ± 0.08 |

* Average $K_d$ value determined from three independent measurements with standard deviation (SD)

**Supplementary Table S3**

**Data collection and refinement statistics of DNA-free HLp, HLp-DNA_1 and HLp-DNA_2.**

Values for the outer shell are given in parentheses. Values for the ellipsoidal completeness are given in square brackets.

| | HLp | HLp-DNA_1 | HLp-DNA_2 |
|---|---|---|---|
| **Data collection** | | | |
| Space group | $P3_121$ | $P4_12_12$ | F222 |
| Cell dimensions | | | |
| *a*, *b*, *c* (Å) | 55.70, 55.70, 58.46 | 67.93, 67.93, 97.90 | 48.77, 91.61, 100.33 |
| α, β, γ (°) | 90, 90, 120 | 90, 90, 90 | 90, 90, 90 |
| Resolution range (Å) | 48.25-1.30 (1.38-1.30) | 43.16-2.10 (2.23-2.10) | 39.56-1.90 (2.05-1.90) |
| Completeness (%) | 99.7 (98.3) | 99.7 (99.5) | 68.4 (19.9) [85.4 (48.4)] |
| Redundancy | 9.43 (5.34) | 5.62 (5.53) | 7.51 (6.44) |
| $\langle I/\sigma(I)\rangle$ | 19.87 (1.14) | 11.09 (1.96) | 13.86 (1.34) |
| $R_{meas}$ | 0.052 (1.40) | 0.081 (0.743) | 0.060 (1.534) |
| **Refinement** | | | |
| No. of reflections, working set | 24883 | 13229 | 5261 |
| No. of reflections, test set | 1303 | 696 | 918 |
| Final $R_{cryst}$ | 0.175 | 0.250 | 0.235 |
| Final $R_{free}$ | 0.211 | 0.280 | 0.288 |
| R.m.s. deviations | | | |
| Bonds (Å) | 0.0046 | 0.0068 | 0.0042 |
| Angles (°) | 1.164 | 1.297 | 1.173 |

**Supplementary Table S4**

**Amino acid residues of the four defined binding sites.**

| Binding site | Residues (Chain ID-Residue ID) |
|---|---|
| **A-site 1** | A-MET26, A-THR27, A-SER28, A-GLY29, B-LYS54, B-ARG55, B-THR56, B-THR57, B-VAL58, B-ARG59, C-MET26, C-THR27, C-SER28, C-GLY29, D-LYS54, D-ARG55, D-THR56, D-THR57, D-VAL58, D-ARG59 |
| **A-site 2** | A-LYS54, A-ARG55, A-THR56, A-THR57, A-VAL58, A-ARG59, B-MET26, B-THR27, B-SER28, B-GLY29, C-LYS54, C-ARG55, C-THR56, C-THR57, C-VAL58, C-ARG59, D-MET26, D-THR27, D-SER28, D-GLY29 |
| **B-site 1** | C-ILE11, C-VAL12, C-ALA-13, C-SER14, C-LYS15, C-LYS17, C-LYS21, D-ILE11, D-VAL12, D-ALA-13, D-SER14, D-LYS15, D-LYS17, D-LYS21 |
| **B-site 2** | A-ILE11, A-VAL12, A-ALA-13, A-SER14, A-LYS15, A-LYS17, A-LYS21, B-ILE11, B-VAL12, B-ALA-13, B-SER14, B-LYS15, B-LYS17, B-LYS21 |

### Supplementary figures

A

>NPDY01000005.1:141450-141799 Leptospira perolatii strain FH1-B-C1 contig_5, whole genome shotgun sequence
CTTCTGAAAATGATTCCTTTTTTGAGGGTATTCTGGAATGGTATCTTTCTAAGTACTGAGTACCTTCTATAAGAACTTACAGAAAAGGAGAAAGAGCTCCCCGGCAA
AAATTGGAT**ATG**GTACAAAAGAAAAAAACTACAGTTAAG**AGGAAG**AGTTCC**ATG**GCTCAAAACGCTGAAAAGGATACCCTCATCGTCGCAAGTAAGGTGAAAGCCTA
TATCAAATCCAAAGGGTTCATGACTTCTGGGGATGCAGTCGATGGTTTGAATGAAAAGTTGTACGCATTAATCGATGATGCGTTAAAGCGCACTGAGTCCAACAAAC
GGACTACGGTTCGCCCAACCGACTTCTAA

>WP_207761458.1 hypothetical protein [Leptospira perolatii]
**M**VQKKKTTVKRKSS**M**AQNAEKDTLIVASKVKAYIKSKGFMTSGDAVDGLNEKLYALIDDALKRTESNKRTTVRPTDF

>PJZ70190.1 hypothetical protein CH360_07900 [Leptospira perolatii]
**M**AQNAEKDTLIVASKVKAYIKSKGFMTSGDAVDGLNEKLYALIDDALKRTESNKRTTVRPTDF

B

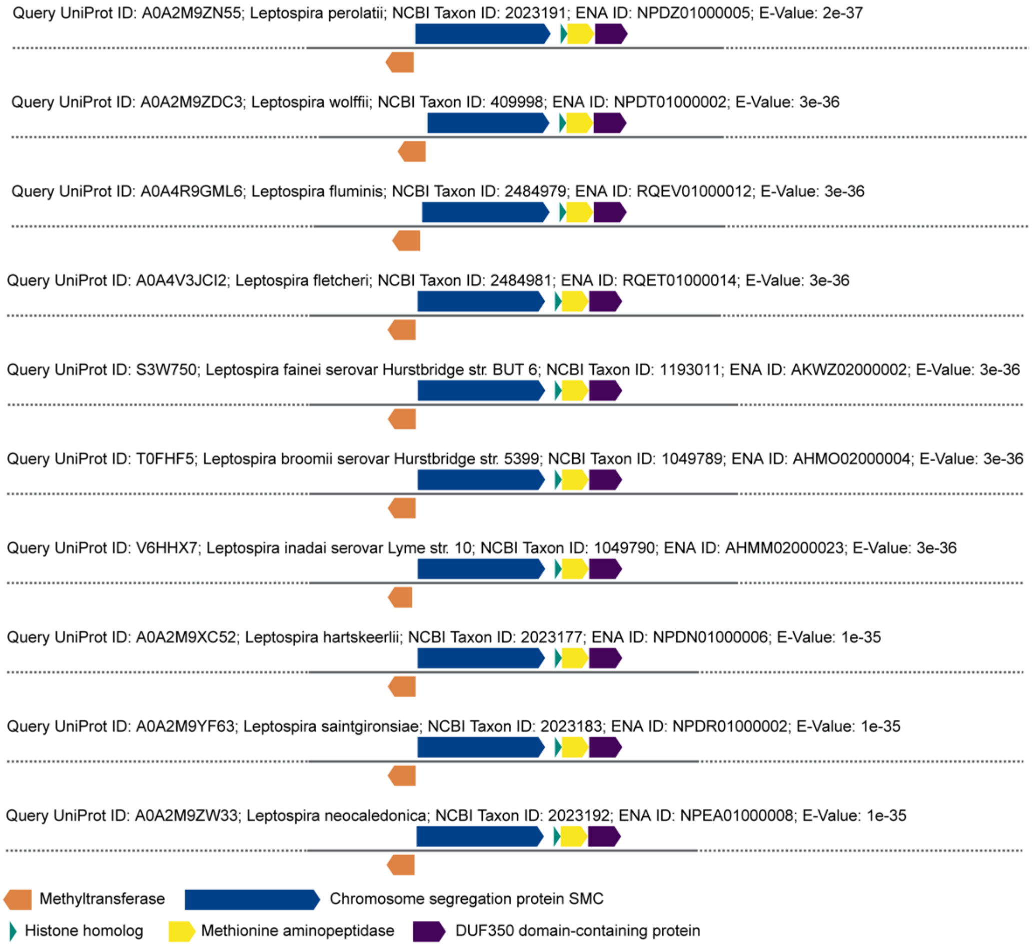


**Supplementary Figure S1**

**DNA and protein sequences of HLp and conserved genomic context across Leptospira strains.**

**A**. Nucleotide and amino acid sequences of HLp. The start codon of the longer *hlp* gene and its corresponding methionine are colored green, while those of the shorter gene variant are highlighted in orange. The Shine-Dalgarno sequence is highlighted in yellow. **B.** Genomic neighborhood diagrams showing HLp and its homologs in various Leptospira species, illustrating conserved synteny and gene context.

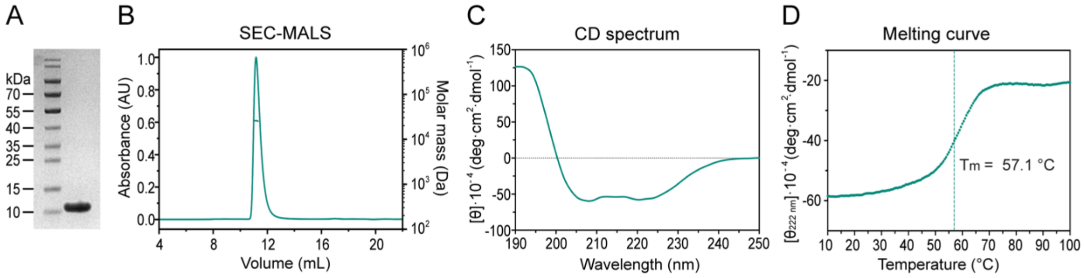


**Supplementary Figure S2**

**Biophysical characterization of HLp in terms of purity, stability, and oligomeric state.**

**A**. SDS-PAGE showing purified HLp. **B.** SEC-MALS analysis of HLp showing its tetrameric state. **C.** Single CD spectrum of HLp. **D.** Thermal melting curve of HLp measured with CD spectroscopy at a wavelength of 222 nm.

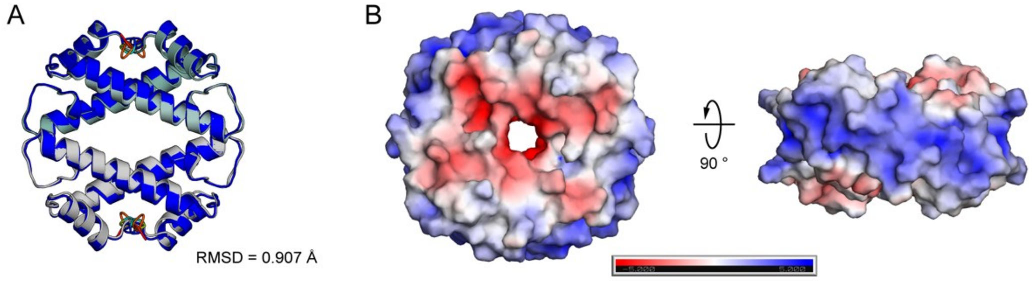


**Supplementary Figure S3**

**Analysis of the HLp crystal structure.**

**A.** Superposition of the crystal structure of the HLp tetramer with its AlphaFold prediction, showing an RMSD of 0.907 Å. **B.** APBS (Advanced Poisson-Boltzmann Solver) electrostatic analysis of the HLp tetrameric structure reveals a continuous, positively charged surface encircling the entire tetramer.

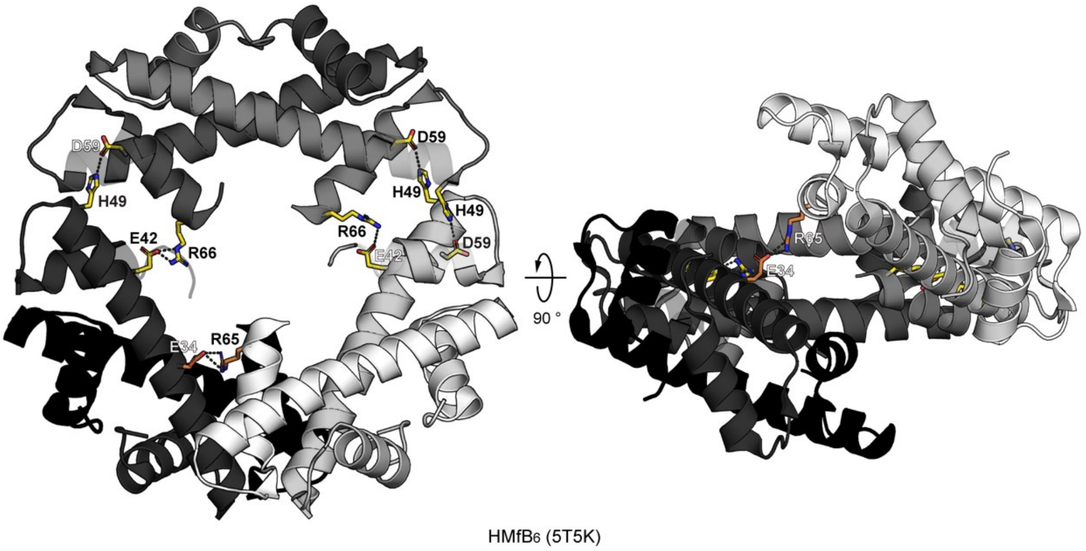


**Supplementary Figure S4**

**Structure of three assembled HMfB dimers.**

Crystal structure of three HMfB dimers in spiral arrangement (PDB: 5T5K) in cartoon representation. Residues involved in oligomerization are shown as sticks and the salt bridges formed between HMfB dimers are indicated as dashed lines.

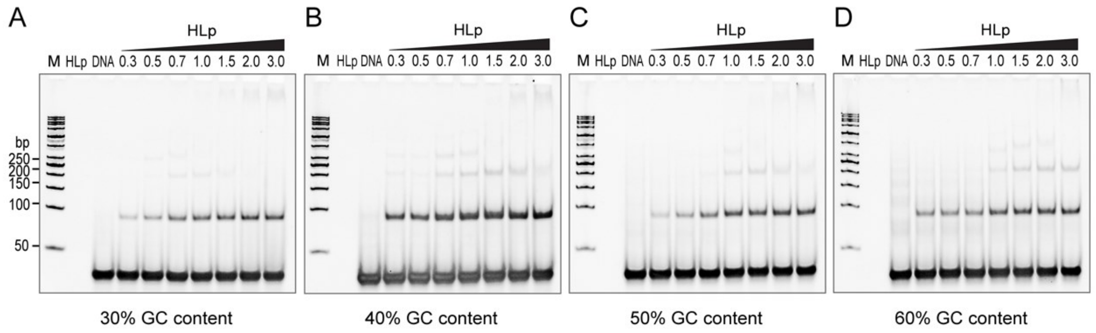


**Supplementary Figure S5**

**Binding of HLp to DNA fragments of variable GC content.**

EMSAs showing binding of HLp to DNA fragments 30-bp-GC30 (**A**), 30-bp-GC40 (**B**), 30-bp-GC50 (**C**), and 30-bp-GC60 (**D**). The molar protein to DNA ratios loaded in lanes 4-10 are indicated.

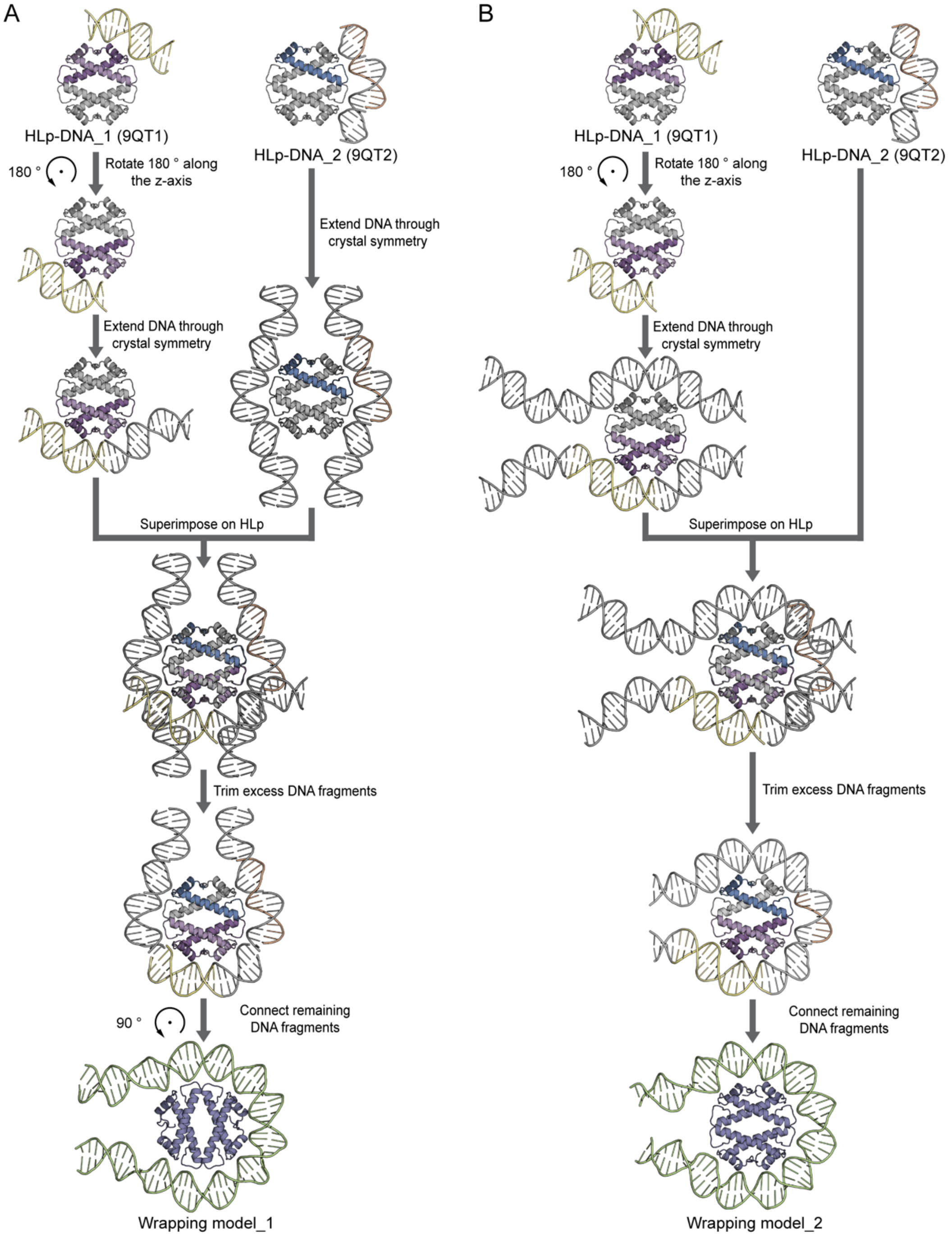


**Supplementary Figure S6**

**The workflow to generate starting models for wrapping mode.**

Workflows, demonstrating how the starting models — wrapping model_1 (A) and wrapping model_2 (B) — were constructed. The crystal structures of HLp-DNA_1 (PDB: 9QT1) and HLp-DNA_2 (PDB: 9QT2) depict the contents of a single asymmetric unit in color, with selected symmetry mates shown in gray. To enhance visualization, the HLp-DNA_1 structure was rotated 180° along the z-axis before extending the DNA fragments through crystal symmetry.

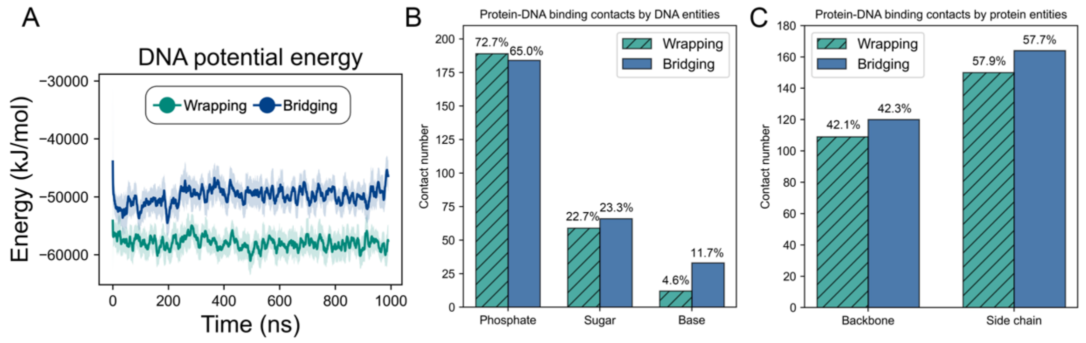


**Supplementary Figure S7**

**DNA potential energy and protein-DNA interactions in the wrapping and the bridging Model.**

**A.** DNA potential energy plotted throughout the simulations. **B.** Average number of protein-DNA atom-atom contacts classified by DNA interaction entities (phosphate groups, sugars, and bases). **C.** Average number of protein-DNA atom-atom contacts classified by protein interaction entities (backbones and side chains). In all panels, the colors green and blue correspond to the wrapping model and bridging model, respectively.

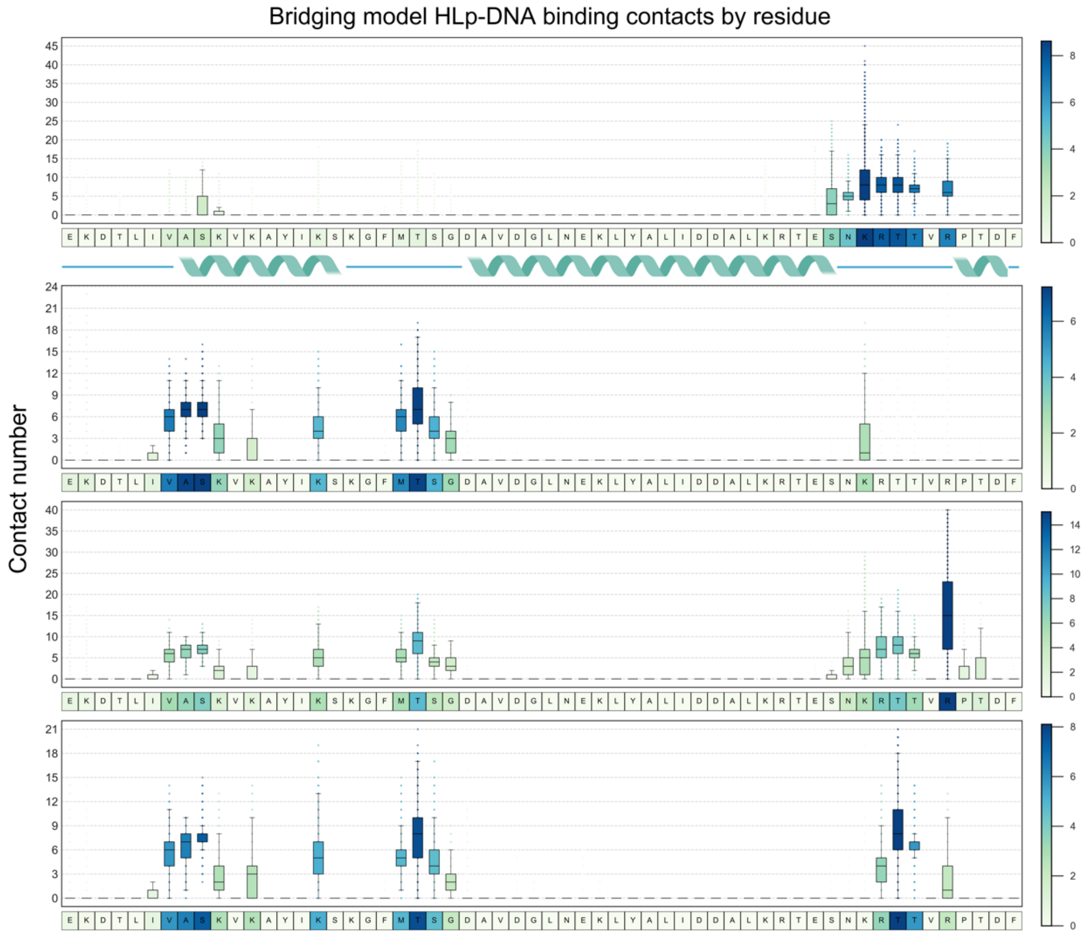


**Supplementary Figure S8**

**Residue-wise binding contact analysis in the bridging model.**

Residue-wise boxplots depicting binding contact numbers for the bridging model. Chains A, B, C, and D are presented sequentially from top to bottom. Darker blue shades indicate higher contact numbers. Below each boxplot, the corresponding amino acid sequence is shown, with a secondary structure diagram provided beneath the first sequence for reference.

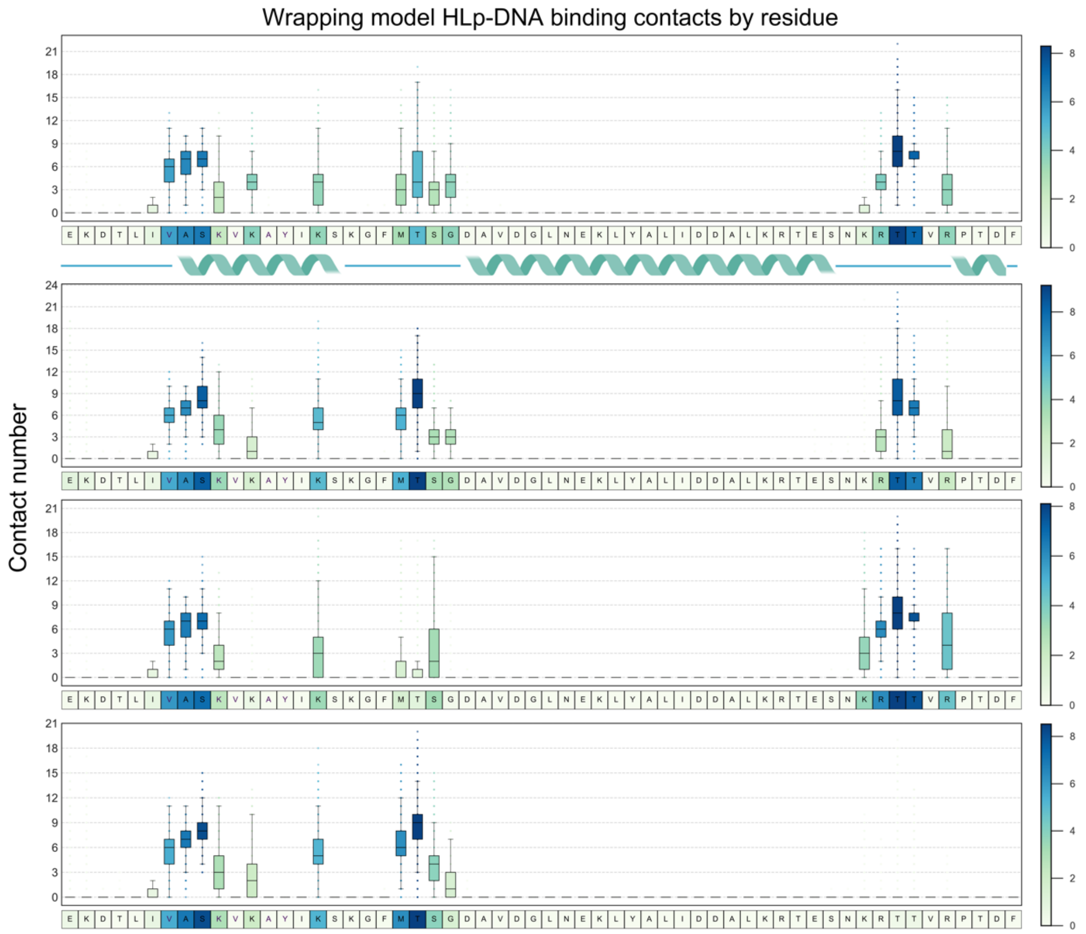


**Supplementary Figure S9**

**Residue-wise binding contact analysis in the wrapping model.**

Residue-wise boxplots depicting binding contact numbers for the wrapping model. Chains A, B, C, and D are presented sequentially from top to bottom. Darker blue shades indicate higher contact numbers. Below each boxplot, the corresponding amino acid sequence is shown, with a secondary structure diagram provided beneath the first sequence for reference.

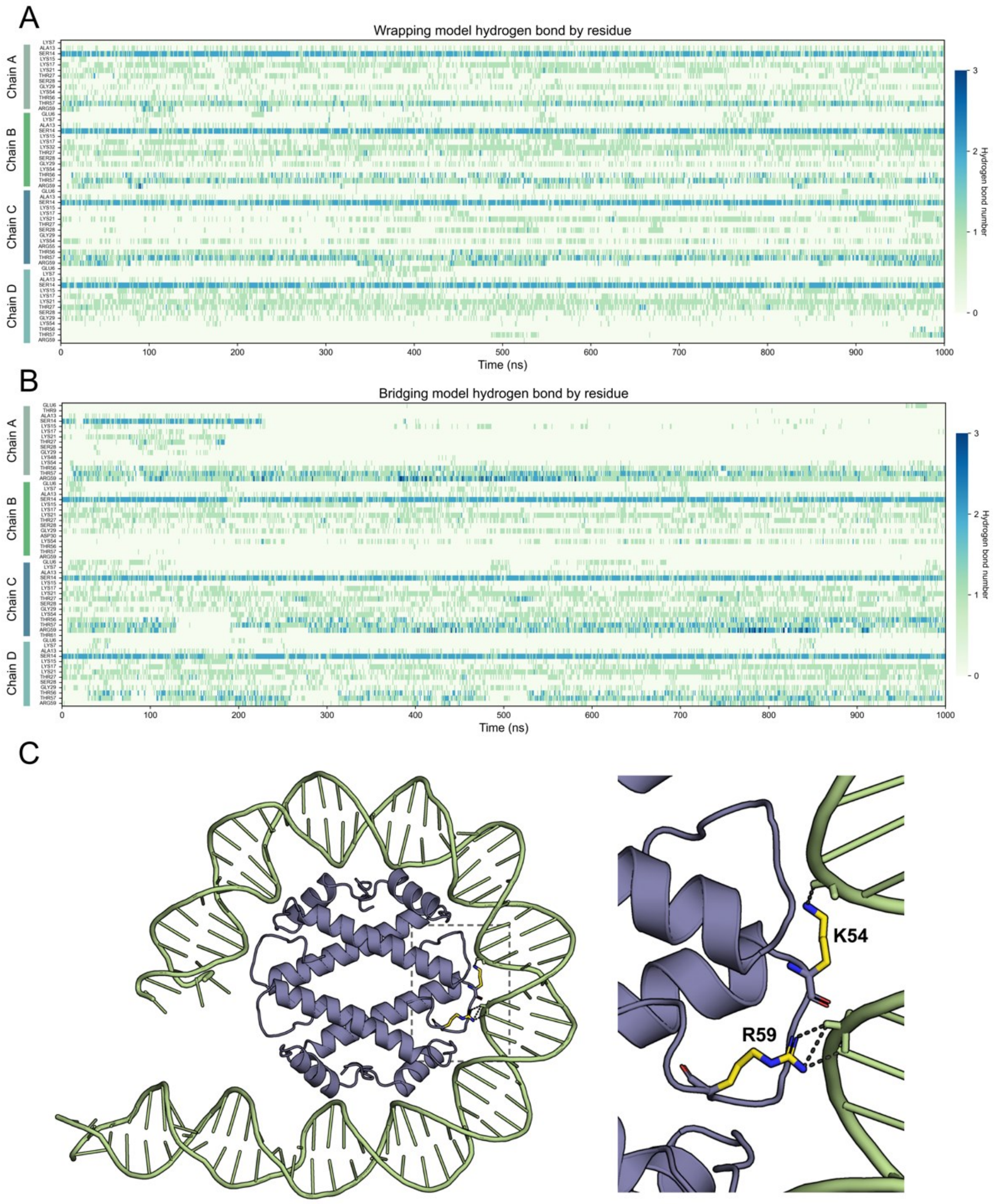


### Supplementary Figure S10

### Protein-DNA hydrogen bond analysis of the wrapping and the bridging models.

Heatmaps depicting the number of protein-DNA hydrogen bonds for each residue in the wrapping (**A**) and bridging model (**B**) throughout the simulation. The chain identifier of each residue is labeled on the left, with darker blue colors indicating a higher number of hydrogen bonds. **C**. Cartoon representation highlighting structural details of residues K54 and R59 in one monomer of the wrapping model, captured at the 150 ns snapshot of the HLp-DNA simulation. Both residues are shown as sticks with yellow-colored side chains, and hydrogen bonds between residues and DNA are represented by dashed lines.

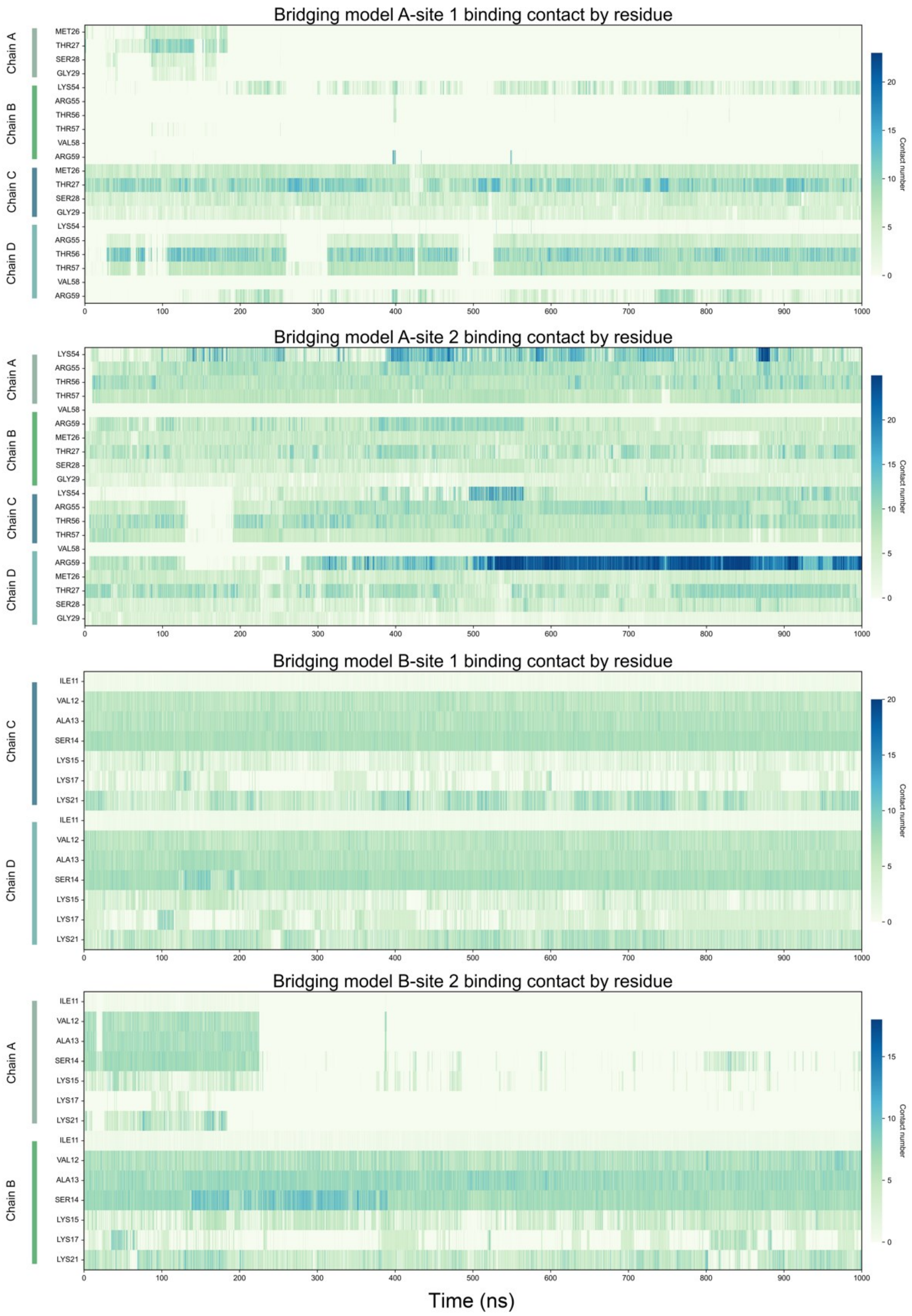


### Supplementary Figure S11

**Protein-DNA contact analysis across binding sites in the bridging model.**

Heatmap illustrating the protein-DNA contact number for each residue across the four binding sites in the bridging model throughout the simulation. Darker blue colors indicate higher contact numbers. The four binding sites — A-site 1, A-site 2, B-site 1, and B-site 2 — are arranged sequentially from top to bottom, with the chain identifier of each residue labeled on the left.

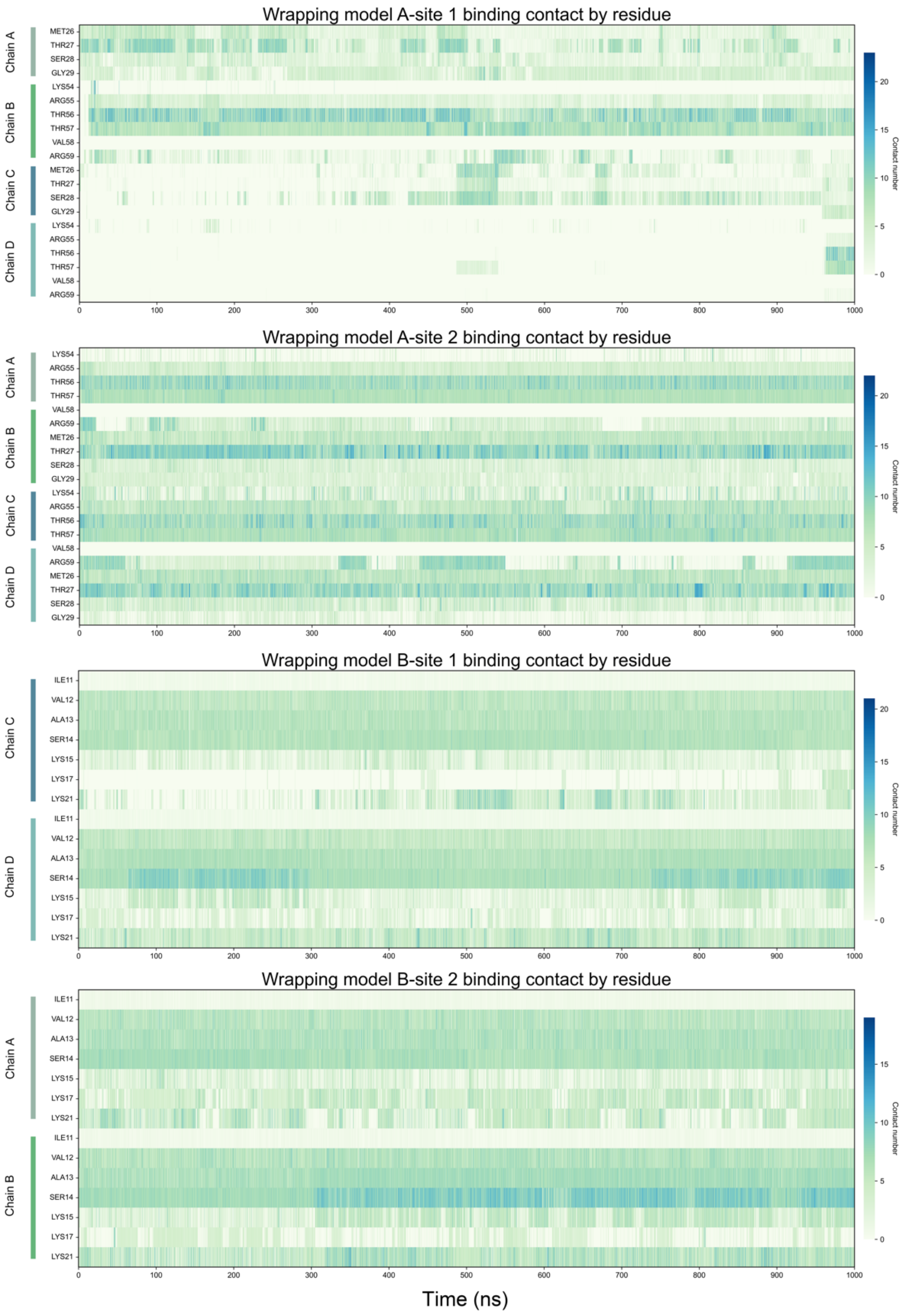


### Supplementary Figure S12

**Protein-DNA contact analysis across binding sites in the wrapping model.**

Heatmap illustrating the protein-DNA contact number for each residue across the four binding sites in the bridging model throughout the simulation. Darker blue colors indicate higher contact numbers. The four binding sites — A-site 1, A-site 2, B-site 1, and B-site 2 — are arranged sequentially from top to bottom, with the chain identifier of each residue labeled on the left.

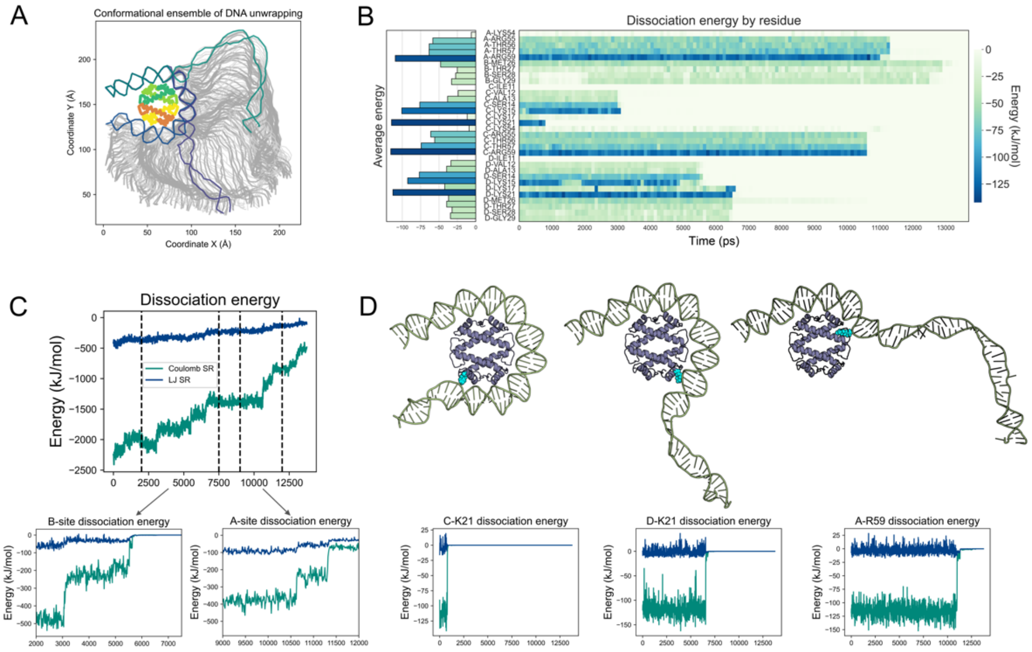


**Supplementary Figure S13**

**DNA unwrapping simulation.**

**A.** This panel shows the conformational ensemble of DNA throughout the simulation. The two ssDNA strands are colored light gray and dark gray, respectively. The HLp tetramer is colored by chains. **B.** Heatmap illustrating the energy profile of each residue involved in A-site 2 and B-site 2 during the unwrapping simulation. Darker blue shades indicate lower energy. The average energy of each residue, excluding frames where the residue is dissociated (with an energy of 0 kJ/mol), is shown as a bar plot on the left. **C.** Energy profile of the HLp-DNA system throughout the simulation, with blue and green lines representing short-range Lennard-Jones (LJ SR) potential energy and short-range Coulomb energy (Coulomb SR), respectively. The dissociation energy of the two binding sites involved in the simulation, B-site 2 and A-site 2, is presented below. **D.** Dissociation energy profiles of three residues with low energy: Chain D-K15, Chain C-R59, and Chain B-K17. These residues are shown as spheres and colored in cyan.

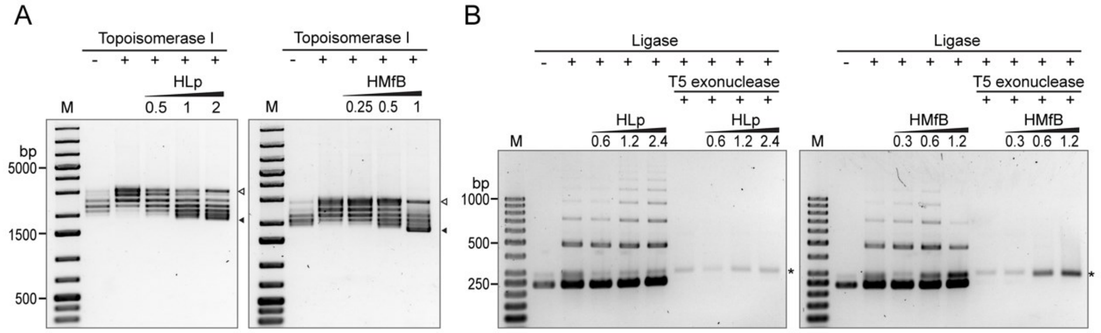


**Supplementary Figure S14**

**HLp binding changes DNA topology.**

**A.** DNA topology assay with relaxed pUC19 plasmid DNA in the presence of HLp and HMfB. Protein to DNA mass ratios are indicated. The bands corresponding to relaxed (△) and supercoiled pUC19 (▲) are labelled. **B.** Ligase-mediated circularization assay with the 240-bp-GC40 DNA and HLp or HMfB. Samples are shown before and after T5 exonuclease digestion. The ratio of protein to DNA mass is labelled. Circularized monomeric DNA is marked (*).

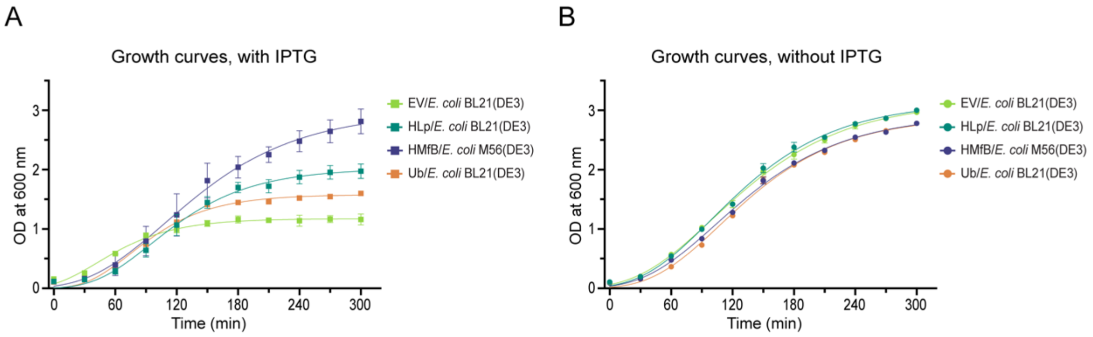


**Supplementary Figure S15**

**Fig. S13 Growth curves of *E. coli* expression strains.**

*E. coli* strains were transformed with empty pET-30a(+) (EV), HLp in pET-30a(+), *Cs*Ub in pET-30a(+), and HMfB in pET-28a(+). Growth curves were recorded in triplicate, with protein expression induced (**A**) and uninduced (**B**) by IPTG.
